# PreMode predicts mode-of-action of missense variants by deep graph representation learning of protein sequence and structural context

**DOI:** 10.1101/2024.02.20.581321

**Authors:** Guojie Zhong, Yige Zhao, Demi Zhuang, Wendy K Chung, Yufeng Shen

## Abstract

Accurate prediction of the functional impact of missense variants is important for disease gene discovery, clinical genetic diagnostics, therapeutic strategies, and protein engineering. Previous efforts have focused on predicting a binary pathogenicity classification, but the functional impact of missense variants is multi-dimensional. Pathogenic missense variants in the same gene may act through different modes of action (i.e., gain/loss-of-function) by affecting different aspects of protein function. They may result in distinct clinical conditions that require different treatments. We developed a new method, PreMode, to perform gene-specific mode-of-action predictions. PreMode models effects of coding sequence variants using SE(3)-equivariant graph neural networks on protein sequences and structures. Using the largest-to-date set of missense variants with known modes of action, we showed that PreMode reached state-of-the-art performance in multiple types of mode-of-action predictions by efficient transfer-learning. Additionally, PreMode’s prediction of G/LoF variants in a kinase is consistent with inactive-active conformation transition energy changes. Finally, we show that PreMode enables efficient study design of deep mutational scans and optimization in protein engineering.

## Main

Accurate and comprehensive prediction of variant effects has been a long-standing fundamental problem in genetics and protein biology. Single amino acid (missense) variants are the most common type of coding variants that contribute to many human diseases and conditions^1–5^. The functional impact of most missense variants remains uncertain. At the molecular level, missense variants in only 40 human genes have been screened in saturated mutagenesis experiments^6^. At the genetic level, only about 2% of clinically observed missense variants are classified as pathogenic or benign, while the majority remain of uncertain clinical significance^7^. Such limitations make it challenging for accurate clinical diagnosis and timely clinical interventions. Furthermore, understanding the functional impact of mutations is important to protein engineering, especially in directed evolution methods, where proteins are iteratively mutated to optimize function or fitness^8^. Such efforts were often limited by the high cost and explosion of sequence space. It remains a challenge to understand and predict the fitness landscape of mutants to reduce the search space and improve the efficiency of engineering^9,10^.

In the past decade, many computational methods have been developed^11–22^ to predict variant effects in a binary manner aiming at distinguishing pathogenic and benign variants. These methods showed that pathogenicity can be predicted by manually encoded or self-learned features based on sequence conservation, protein structures, and population allele frequency. Moreover, recently developed methods based on protein language models, leveraging Transformer architectures and self-supervised training on billions of protein sequences from UniProt^23^, have demonstrated their capability to serve as versatile predictors of various protein features^24–26^. The embeddings from these models can offer zero-shot predictive potential for variant pathogenicity^27,28^. While those methods are helpful in genetic analyses, pathogenicity does not capture the complexity of functional and genetic effects of variants. For example, gain of function variants in *SCN2A* lead to infantile epileptic encephalopathy^29,30^ while loss of function variants in the same gene lead to autism or intellectual disability^29,30^. Such limitation reduced the utility of the methods in genetic analysis and clinical applications.

We use “mode-of-action” as a generic term to encapsulate the multi-dimensional molecular and genetic mechanisms through which pathogenic variants impact protein functionality and increase the risk of diseases, respectively. More specifically, at molecular level, pathogenic variants can change the biochemical properties of a protein in different ways. For example, decreasing/maintaining protein stability^31,32^, enzymatic activity^32,33^, regulatory functions, and interaction^34,35^. At genetics level, variants are often categorized into two major types, gain or loss of function (G/LoF). GoF variants encompass alterations that perturb the protein from its normal functions via increased or novel activities^36,37^. GoF variants are often found to be driver mutations in oncogenes^38^. LoF variants damage protein function via decreased activities, which are often found in tumor suppressors in cancer^39^ and other genetic diseases^40^. Gain and loss of function variants usually result in markedly different clinical phenotypes^36,41–44^, necessitating entirely distinct therapeutic approaches^29,35,36,45^.

While numerous methods have demonstrated the potential to predict pathogenicity on a genome-wide scale, the effort in G/LoF prediction has been limited. Stein et al attempted to predict genome-wide G/LoF variants via assembly of human curated features^46^. However, we note that mode-of-action centers around how a variant disrupts the normal function of a protein. Given the inherent diversity of protein functions, attempting to define a universally applicable predictive task for all G/LoF variants across all proteins could lead to conceptual ambiguity. Therefore, we propose that such predictive tasks should be defined within the context of individual proteins or protein families that share similar functions. The main challenge is the limited availability of data for most genes and protein families.

We developed a new method, PreMode (Pretrained Model for Predicting Mode-of-action), to address these challenges with deep learning models through genome-wide pretrain and protein-specific transfer learning. PreMode is designed to capture the variant impact on protein function with regard to its structural properties and evolutional information. We built PreMode with SE(3)-equivariant graph attention transformers, utilizing protein language model embeddings^25^ and protein structures^24^ as inputs. We curated the largest-to-date labeled missense variants with mode-of-action annotations from clinical databases, genetic inference, and experimental assays. We applied PreMode to mode-of-action predictions of 17 genes. PreMode reached state-of-the-art performance at mode-of-action predictions compared to existing models. We further demonstrate PreMode’s practical utility in both improving data analysis in deep mutational scan experiments and assisting protein engineering by significantly reducing the size of mutants for screening via active learning.

## Result

### Overview

We proposed a framework for predicting the mode-of-action at the molecular level and genetic level. Molecularly, the effect of a missense variant is about the change in biochemical properties of a protein, such as enzyme activity, stability, and the regulatory processes upon protein-protein interactions (**Figure 1a**). These changes can be measured by deep mutational scan experiments (**Figure 1a**). Genetically, the overall outcome of molecular effects results in different types of missense variants. One common categorization is “loss of function” (LoF) and “gain of function” (GoF) variants (**Figure 1a**). To conceptualize this framework to variant effect prediction models, we introduced two parameters: “distance from wild type” (denoted as ‘*r*’) and the “direction of change” (notated as ‘θ’). The distance parameter distinguishes between pathogenic and benign variants (**Figure 1a**) and is shared across all genes. The direction parameter takes on different meanings both molecularly and genetically within various genes. Therefore, we proposed that a mode-of-action predictor would make separate predictions of *r* and θ utilizing different datasets. It would first learn *r* prediction using labeled pathogenic and benign variants for all genes, just like conventional variant effect predictors, then learn θ prediction using protein or protein family specific datasets via transfer learning (**Figure 1b**).

**Figure 1.**
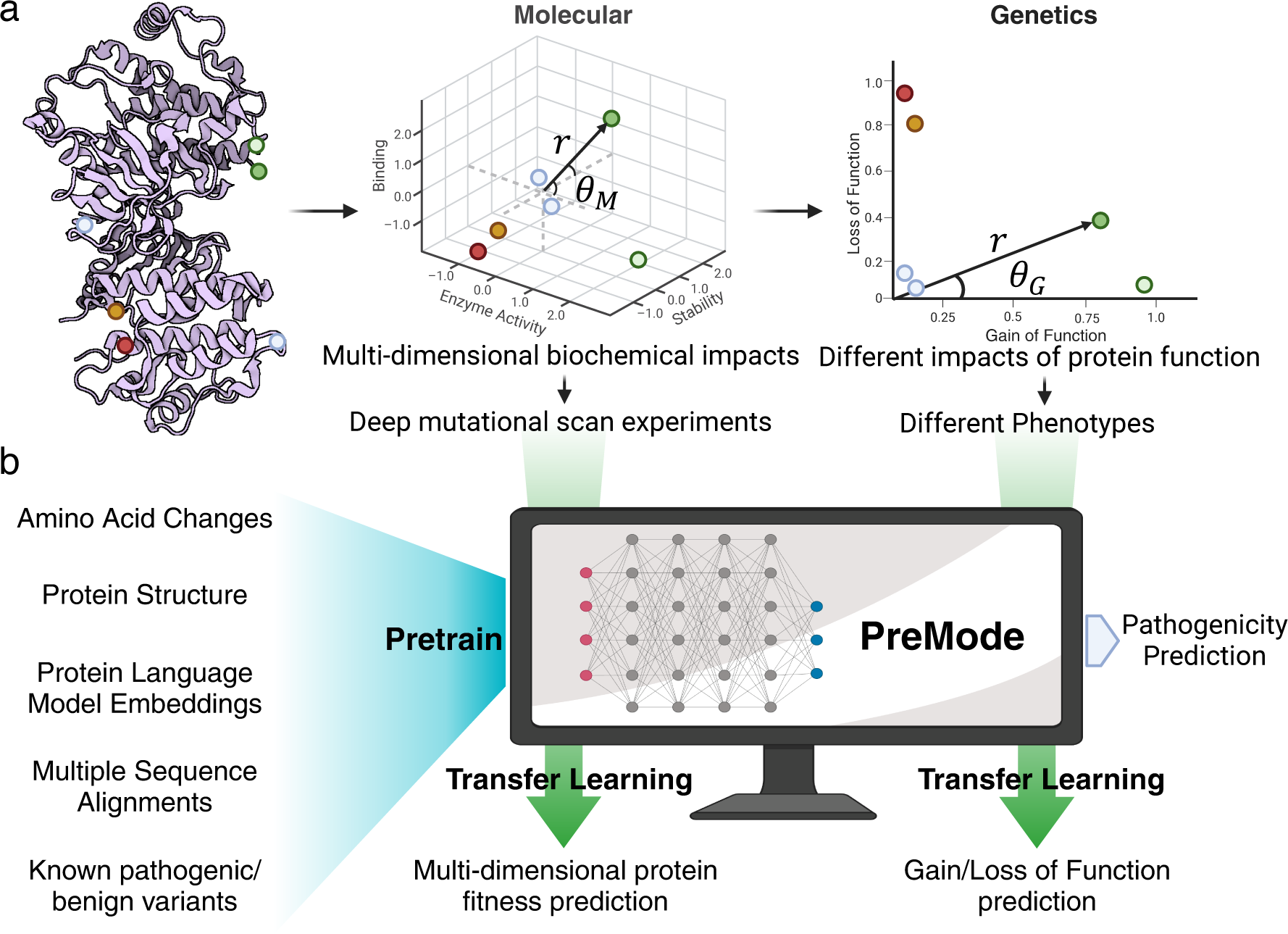
Mode-of-action definition and PreMode framework. A) Mode-of-action definition at molecular level and genetic levels. B) PreMode framework. Blue arrows show the data flow at pretrain stage for pathogenicity prediction, green arrows show transfer-learning stage for mode-of-action prediction.

### Curation and characterization of mode-of-action labeled missense variants

We curated the largest-to-date mode-of-action labeled missense variants datasets annotated at both molecular and genetic levels, including 41,081 missense variants in eight genes with multi-dimensional measurements of different biochemical properties by deep mutational scan experiments curated from MAVEDB^6^, 2043 gain- and 7889 loss-of-function missense variants in ∼1300 genes. The gain and loss of function-labeled variants were collected from literature searches^36,44,47,48^, cancer hotspots^49–51^ and published databases^52^ (**Methods**).

We first investigated global properties of GoF and LoF variants using the curated data set. GoF variants are more likely to be located in regions with lower AlphaFold2 prediction confidence (pLDDT) than LoF variants and are more likely to be on protein surfaces (**Figure 2a**). In contrast, LoF variants have an overall bigger impact on protein folding energy than GoF variants (Figure 2a), although both types of variants confer a greater folding energy change to protein folding than benign variants (**Figure 2a**). As expected, both LoF and GoF variants are more likely to be in conserved regions than non-conserved regions, as shown in distribution of conservation represented by entropy of amino acid frequencies across species in multiple sequence alignments (MSA) (**Figure 2a**), while benign variants are mostly located in non-conserved regions (**Figure 2a**). Additionally, GoF variants in general are more likely to be located in disordered regions without specific secondary structures than LoF variants (**Figure 2b**).

**Figure 2.**
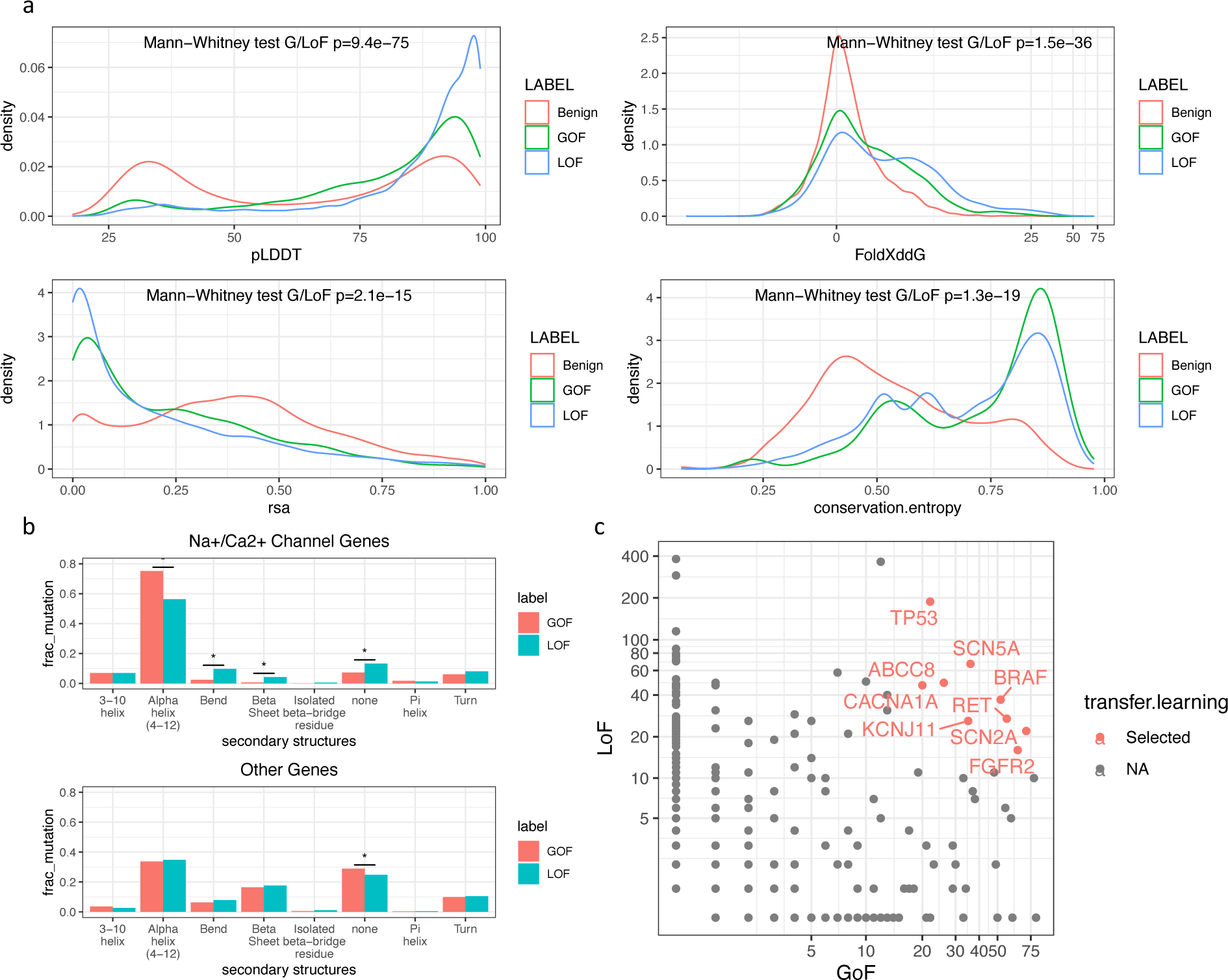
Characterization of gain-/loss-of-function variants. A) Genome wide comparison of gain- and loss-of-function variant sites’ pLDDT (AlphaFold2 prediction confidence), protein folding energy change, relevant solvent accessibility, and conservation. B) Protein family wise comparison of gain- and loss-of-function variants enriched secondary structures. C). Gene wise comparison of gain- and loss-of-function missense variants numbers.

However, we note this pattern is different across protein families. For example, in Na+/Ca2+ channel genes, GoF variants are more enriched in alpha helixes that are critical for ion transport and selectivity domains than LoF variants (**Figure 2b**). Finally, the number of GoF and LoF variants are not evenly distributed across genes, with only a few of the genes having more than 15 GoF and LoF variants (**Figure 2c**). Overall, those results showed that protein structure, energy, and evolutionary features could help predict G/LoF variants while underscoring the necessity for the development of protein- and protein family-specific predictive models using limited data.

### A deep learning model for mode-of-action predictions

We developed PreMode, a model pre-trained on pathogenicity prediction task and optimized for transfer learning to mode-of-action prediction tasks. PreMode takes input features derived from amino acid biochemical properties, protein contexts, and cross-species conservation. PreMode models protein 3D context structure with SE(3)-equivariant graph neural networks (**Figure 3a**). PreMode was designed to not only capture the relative importance between residues by taking both backbone torsion angles and side chain directions into consideration, but also maintaining awareness of geometric equivariance so that rotation of the atom coordinates does not affect the predictions. PreMode’s SE(3)-equivariant learning ability was achieved by using a graph representation of protein 3D structures, where each residue was represented as nodes with features that explicitly represent local biochemical properties and evolutionary conservation including secondary structures^53,54^, pLDDT^24^, amino acid frequencies in MSA^55^, and relative coordinates of all atoms in sidechain with respect to alpha carbons (**Methods**). We also included protein sequence language model (ESM2) embeddings^25^ into node embeddings, which implicitly capture similar structural and evolution information. Such implicit representation could serve as a compensation of possible missing information limited by secondary structure annotation or MSA generation algorithms. For each edge connecting two residues, the features include Euclidean vector of two corresponding beta-carbons (for glycine we use alpha-carbon instead) to encapsulate static contacts. Additionally, we incorporate co-evolutional strength calculated from MSA and contact strength predictions from MSA transformer^56^ to represent potential dynamic contacts^57^ or inter-homopolymer contacts that are missed in a static structure (**Methods**). PreMode applies SE(3)-equivariant mechanisms on edge features and node features first through a star-graph that connects variant site with all other residues to capture the direct impacts of amino acid alterations on other residues, then through the k-nearest neighbor (KNN)-graph that connects each residue with its closest neighbor residues to capture the second order impacts, finally through a star-graph module to aggregate the impacts (**Supplementary Figure 1**).

**Figure 3.**
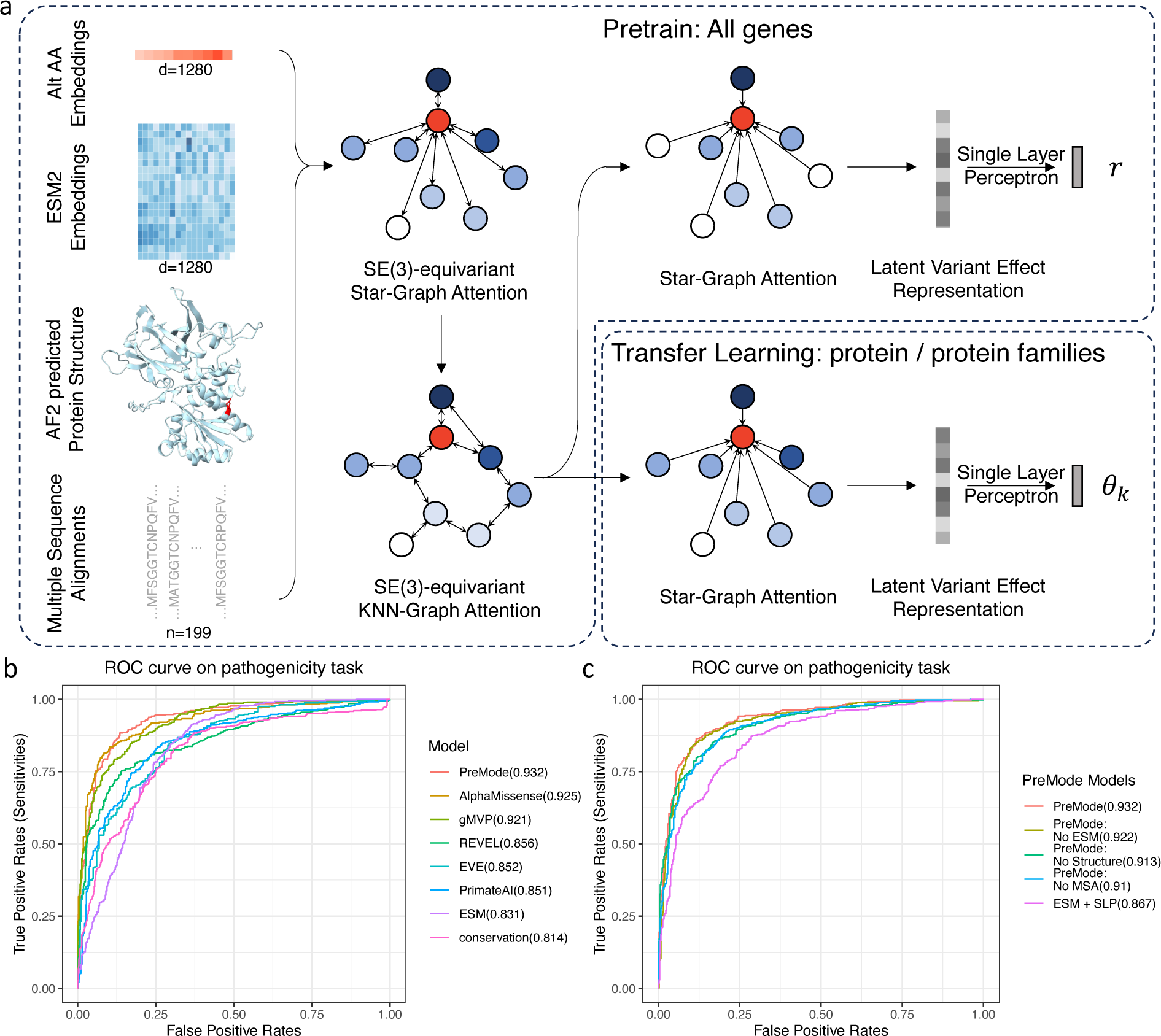
PreMode, a pretrained model for mode-of-action predictions. A) PreMode framework. PreMode takes amino acid changes, protein language model embeddings, alphafold2 predicted structures, multiple sequence alignments as inputs and outputs two parameters. PreMode first predicts pathogenicity (*r*) for all genes during pretrain, next predicts the mode-of-action parameters (θ) via transfer learning. B) AUROC of PreMode and other methods in pathogenicity prediction task. C) AUROC of PreMode in ablation analysis.

We pretrained PreMode using labeled pathogenicity data to let the model learn general representation of the variant effects. We collected 83,844 benign variants from ClinVar and PrimateAI^17^ in 13,748 genes, and 64,480 likely pathogenic variants from ClinVar with at least one star confidence and HGMD in 3703 genes. We randomly selected 5% of the variants as a validation dataset and trained 20 epochs on the rest of the training data until validation loss stopped dropping (**Supplementary Figure 2**). While predicting pathogenicity is not the designed goal of PreMode, we can still use pathogenicity prediction performance to investigate the contribution from various components of the model. As PreMode was pre-trained on human-curated ClinVar data, using variants from the same resource as testing data can result in inflated performances. Instead, we used independent testing data for which the pathogenicity label was entirely based on statistical evidence, that is, 533 pathogenic missense variants in cancer hotspots from cBioportal^49,51,58^ and same number of benign variants in the same genes randomly selected from common variants in primates^17^.

PreMode reached similar levels of performance as existing methods including AlphaMissense^20^ and gMVP^19^ on the testing dataset with AUROC (area under recall receiver operating characteristic curve) of 0.932 (**Figure 3b**). We performed ablation analysis to assess the contribution of language model embeddings, structural information and MSA information to the prediction. Replacing ESM2 embeddings with one-hot encodings of amino acids resulted in a drop of AUROC to 0.922. Similarly, removing the MSA will drop the AUROC to 0.91 (**Figure 3c**). Removing the structure module will drop the performance slightly to 0.913 (**Figure 3c**). This showed that ESM2 embeddings, MSA module and SE(3)-equivariant module on AlphaFold2 predicted protein structures together provide non-redundant information for pathogenicity prediction.

### PreMode reaches state-of-the-art in molecular mode-of-action predictions and facilitates interpretation of deep mutational scan experiments

We first investigated the utility of PreMode in predicting modes of action at the molecular level. We obtained DMS data on eight genes (*PTEN, SNCA, CCR5, CXCR4, NUDT15, CYP2C9, GCK, ASPA*) from MAVEDB^6^ with multiple assays of different biochemical properties. These assays broadly target two aspects of protein function, the stability and enzymatic activity. These two types of functional readout are moderately correlated as reduced protein stability often directly or indirectly affects function (**Supplementary Figure 3**).

We split the data of each gene into 80% of training and 20% of testing data five times under different seeds and ran PreMode on each of them via transfer learning. While most other methods for pathogenicity prediction does not provide model weights for us to do transfer learning, we compared PreMode against four models (Augmented ESM1b, Augmented EVmutation, Augmented Unirep, Augmented EVE, **Methods**) with top transfer learning performances described in Hsu, et al.^59^, and a baseline model utilizing ESM2 embeddings and a single layer perceptron (SLP, **Methods**) as approximated transfer learning with ESM2. PreMode outperformed all methods with higher spearman correlation (**Figure 4a**) on all the assays of 8 genes. Overall, after transfer learning, PreMode is able to predict both the multi-dimensional protein stability and functional fitness with a spearman correlation of 0.6 with experimental results, better than all other methods (**Figure 4b**). Furthermore, we investigated the multi-dimensional transfer learning ability of PreMode under smaller sample sizes, where we randomly subsampled the training data and compared the performances in same testing dataset. We found that PreMode is better than all other methods in transfer learning with ≥20% training data, while all methods had similar performance under smaller data sizes (**Figure 4c**). Overall, PreMode is able to accurately predict variant effects of all missense variants with around 40% (∼2000) of variants measured inside one gene, after which increasing the number of data points will have minimal improvement on the performance in the testing dataset (**Figure 4c**).

**Figure 4.**
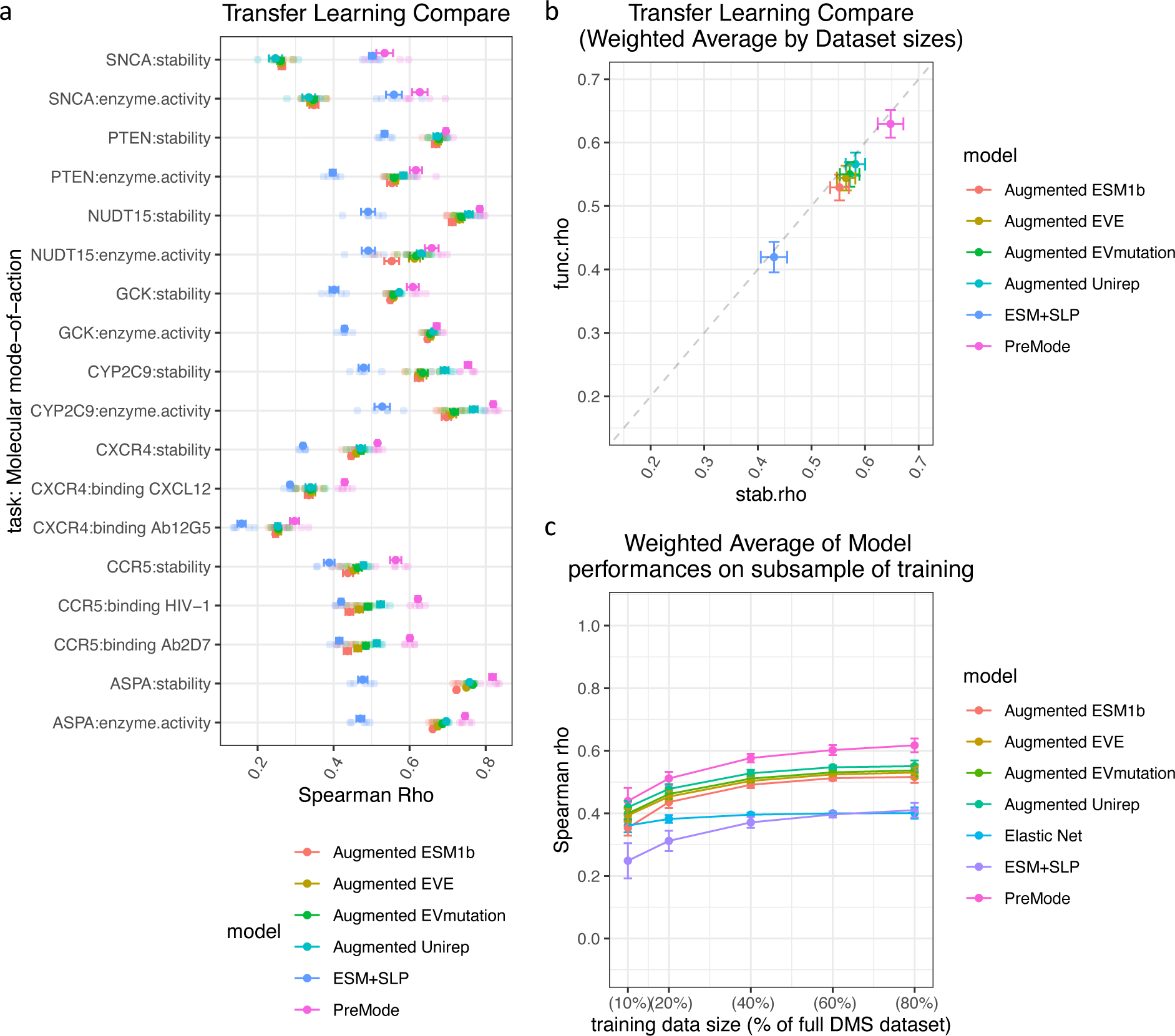
PreMode and other methods’ performances in molecular mode-of-action prediction tasks. A) Spearman correlations of PreMode in multiplexed deep mutational scan experiments of 8 genes compared to other methods. B) Comparison of weighted average of spearman correlations across experiments based on dataset sizes. C) Comparison of weighted average of spearman correlations of PreMode and other methods under sub-sampled data.

We further investigated the utility of PreMode to improve the analysis of experimental readouts in two applications. First, we hypothesized that PreMode scores could indicate abnormal measurements in each experiment by transfer learning as it had implicitly modeled the fitness of variants in all proteins during pretraining. We used the stability DMS experiment^31^ of *PTEN* as an example and trained PreMode on one of the eight biological replicates. We then compared the differences between PreMode’s predictions and the experimental readouts. We showed that this difference value is highly correlated to the difference between the readout of the single biological replicate and average readouts in all experiments (**Supplementary Figure 5a**). The experimental readouts with large deviation from PreMode’s predictions are more likely to be abnormal measurements (**Supplementary Figure 5b**). Next, we hypothesized that models trained on stability in a subset of genes are generalizable to other genes. We applied PreMode to the largest stability measurement experiments in MAVEDB across >30 genes. We trained PreMode on 80% of the data and tested on the other 20% of variants in completely different genes from training. PreMode outperformed all other methods (**Supplementary Figure 6**).

### PreMode is state-of-the-art in genetic mode-of-action predictions

We grouped the gain / loss of function variants dataset by genes and only kept those with ≥15 G/LoF variants (**Figure 2c**, **Methods**). We performed transfer learning on the selected genes using the pretrained model parameters as initial weights (**Methods**). We note there are two reasons to perform transfer-learning in individual genes rather than training a general model across all genes. First, G/LoF mechanisms are intrinsically different across genes, as they have different functions. Second, the number of G/LoF variants are extremely unbalanced across genes. A deep neural network model with transfer-learning across genes will potentially reach a local minimum where gene-properties dominate its predictions that are better at predicting likely G/LoF genes but do not distinguish G/LoF variants in the same gene (**Supplementary Figure 7**).

For each gene, we randomly split the gain and loss of function variants into training and testing. The total amount of data for training and testing in each gene is shown in **Supplementary Table 1**. There were nine genes (*ABCC8, BRAF, CACNA1A, FGFR2, KCNJ11, RET, SCN2A, SCN5A, TP53*; **Figure 2c**) in total. We compared PreMode against several baseline methods trained and tested on the same training/testing datasets. Overall, PreMode performed better in all the genes than baseline methods. It reached average AUC of 0.8∼0.9 in genes *RET, KCNJ11*, *CACNA1A* and *BRAF,* average AUCs of 0.7∼0.8 in *SCN5A, SCN2A, TP53,* and *ABCC8* (**Figure 5a**). PreMode is better than low-capacity models such as random forest using manually curated biochemical features and conservation information as input (**Figure 5a, Methods**). Pretrained PreMode is better than non-pretrained model for all genes except *FGFR2*, where all three models have average AUC lower than 0.6 (**Figure 5a**). We also compared our model to LoGoFunc^46^, a method trained on G/LoF variants across genes. As the training and testing split information were not available for LoGoFunc, we re-split the training and testing data and completely removed the data curated from LoGoFunc in testing datasets for fair comparison. PreMode has similar performance as LoGoFunc in *CACNA1A* but better in all other genes (**Figure 5b**). We next compared our model to FunCion^36^, a method trained only on voltage-gated sodium and calcium channels (encoded by *SCNxA* and *CACNA1x* family genes) with manually curated sequence and structural features. We found that PreMode is better than FunCion in all three genes *SCN2A*, *SCN5A, CACNA1A*, and all ion channel genes when using the same training testing data split (**Figure 5c**).

**Figure 5.**
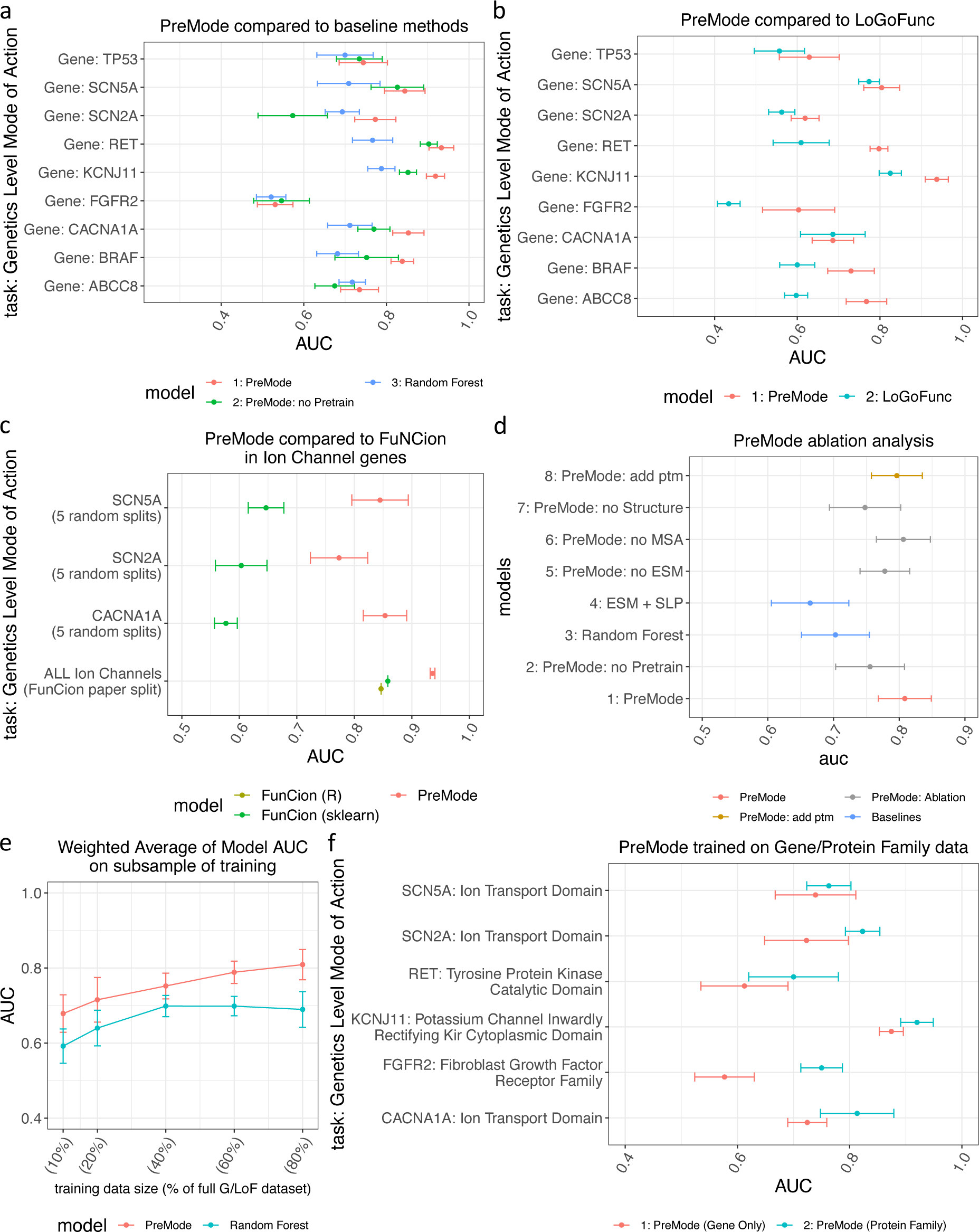
PreMode performances in genetic mode-of-action prediction tasks. A) Performances of PreMode and baseline methods on gain- and loss-of-function predictions in 9 genes compared to random forest and PreMode without pretrain. B) Performances of PreMode and LoGoFunc in 9 genes, training/testing data was split by presence in LoGoFunc data or not (**Methods**). C) Comparison of PreMode and FuNCion in three ion channel genes, training/testing data was split randomly or same as in their paper (**Methods**). D) Ablation analysis of PreMode on gain- and loss-of-function predictions in 9 genes, for each model, AUC in 9 genes were weighted summed by the dataset sizes (**Methods**). E) Few-shot transfer learning of PreMode and random forest method on subsampled training data, AUC in 9 genes were weighted summed by dataset sizes. F) PreMode performances when trained with G/LoF variants from the same gene (marked as “Gene Only”) or G/LoF data from same domain across different genes (marked as “Protein Family”) and tested on the same testing dataset.

We next did ablation analysis on PreMode to identify the important features for Gain/Loss of function predictions. Removing ESM, MSA or protein structure information decreased PreMode’s overall performances in 9 genes (**Figure 5d, Methods**). Removing structural input decreased the performance in all genes except for *FGFR2* (**Supplementary Figure 8a**). We noticed that both the LoF variants and GoF variants in that gene were located in slightly lower pLDDT region (**Supplementary Figure 8c**). Adding post-translational-modification (PTM) information (predicted by PhosphoSitePlus^60^) into the model input can slightly improve the predictions in *TP53, SCN5A,* and *FGFR2.* (**Supplementary Figure 8b**). In these genes, the LoF variants are closer to the PTM sites than GoF variants while the trend is not significant in the other genes (**Supplementary Figure 8d**). Overall, PreMode reached highest performance in the weighted sum of AUC across genes, followed by ablation models and baseline methods (**Figure 5d**).

To be effective in genetic analysis, a machine learning method should be able to perform transfer learning with limited amount of data, as in most of the genes there are fewer than 10 known G/LoF variants (**Figure 2c**). We therefore examined PreMode’s performance with subsampled training data and compared with baseline methods like random forest. We found PreMode’s overall performances in 9 genes were better than random forest even if we down sample the training data to 10% of whole dataset (**Figure 5e**). Furthermore, random forest’s overall performance stopped increasing after we increase the training data size to 40% of whole dataset, while PreMode’s performance is not saturated (**Figure 5e**).

Next, we hypothesized that the gain and loss of function mechanisms are similar in the same domain across genes, and can use this to further improve the GoF/LoF predictions in each gene. We split the data in each gene by domains and only selected the domains with ≥15 G/LoF variants for evaluation, including ion transport domains, tyrosine protein kinase domains and growth factor receptor domains. We performed PreMode transfer learning within the domain using either gene-specific data or data across genes while tested on the same data within one gene. We observed increased performance in all of the domains when using data across genes rather than using the gene alone (**Figure 5f**).

Overall, those result suggested that PreMode is the state-of-art method in G/LoF prediction even with limited data. For genes with small sizes of labeled G/LoF variants data, PreMode could utilize the information from G/LoF variants data in same protein family to further increase performances.

### PreMode predicted mode-of-action landscapes in individual proteins through in silico saturated mutagenesis

We applied PreMode to infer the mode-of-action of all possible variants in *BRAF, TP53, PTEN, RET*, and *KCNJ11*. We averaged the prediction scores of all five models that trained on different training/testing split and applied to all the other variants in corresponding gene.

In *BRAF*, PreMode identified two regions enriched with GoF variants. One is on a N-terminal phorbol-ester/DAG-type zinc-finger domain and the other is on the kinase domain which includes the well-known V600 GoF position^61,62^ (**Figure 6a**). The variants on the two domains act through different gain/loss of function mechanisms. The phorbol-ester/DAG-type zinc-finger domain auto-inhibits BRAF activity through binding with 14-3-3^63^ while also cooperating with Ras-binding domain (RBD) to bind with Ras and activate BRAF activity^64^. To interpret PreMode’s predictions, we obtained B-Raf/14-3-3 and B-Raf/K-Ras binding structures from PDB (7MFD) and AlphaFold2 (Colabfold^65^ implementation) predictions, respectively (**Supplementary Figure 9a**). We then calculated the energy change on both structures upon mutations using FoldX^66^. We found that the GoF variants predicted by PreMode only destabilize B-Raf binding to 14-3-3 and maintain the binding stability to K-Ras at similar level as benign variants (**Figure 6b**). The ddG landscape suggested that the GoF variants in this region mostly act by abolishing its inhibitory regulation^67^. We further investigated the G/LoF variants predicted by PreMode in the kinase domain. The kinase domain has both active (PDB: 4MNE) and inactive (PDB: 4EHE) conformations with large and small enzyme pocket sizes, respectively^68^ (**Supplementary Figure 9b**). The FoldX ddG results were consistent with previous findings that GoF variants V600E/D can destabilize the inactive state and stabilize the active state^69^, while the predicted LoF variants destabilize both conformations (**Figure 6c**). Similarly, we found that predicted LoF variants can destabilize the complex of BRAF-MEK1 while GoF variants maintains the stability of BRAF-MEK1 complex at similar level of benign variants (**Figure 6d, Supplementary Figure 9c**).

**Figure 6.**
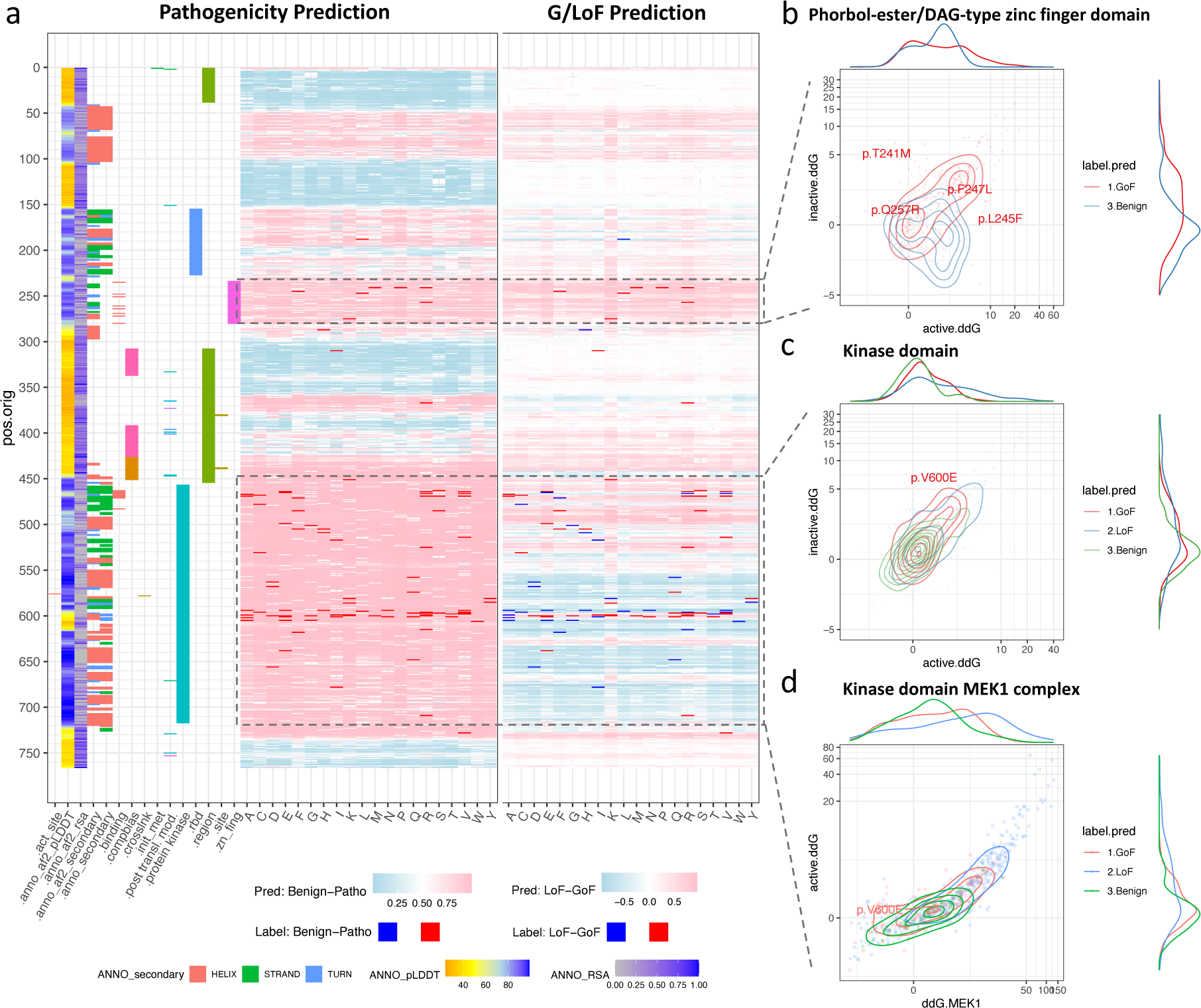
PreMode in-silico mutagenesis mode-of-action predictions on *BRAF*. A) PreMode predictions visualized with BRAF protein domains, x-axis are annotations and amino acid changes, y-axis are amino acid positions. The first few columns indicate Alphafold2 pLDDT, relevant solvent accessibility, secondary structures, protein domains, respectively. The left panel showed PreMode’s pathogenicity predictions and labels used in training, pink indicates predicted pathogenic, light blue indicates predicted benign, red indicates pathogenic and blue indicates benign in training data, white indicates reference amino acid. The right panel showed PreMode’s G/LoF predictions and labels used in training, pink indicates predicted GoF, light blue indicates predicted LoF, white indicates predicted benign. Red indicates GoF and blue indicates LoF in training data. B) Folding energy change upon mutations of Braf phorbol-ester/DAG-type zinc finger domain on the inactive state (Braf-14-3-3 binding, y-axis) versus active state (Braf-Kras binding, x-axis), colored by PreMode predictions, a few known cancer driver mutations were labeled. C) Folding energy change upon mutations of Braf kinase domain on the inactive state (y-axis) versus active state (x-axis), colored by PreMode predictions, a few known cancer driver mutations were labeled. D). Folding energy change upon mutations of BRAF-MEK1 complex. X-axis showed the ddG on BRAF-MEK1 complex, y-axis showed the ddG on BRAF active conformation alone (left panel, PDB: 4MNE), colored by PreMode predictions.

*TP53* is a tumor suppressor gene, and most of the pathogenic variants act through LoF. However, there are a few regions enriched with GoF variants identified by PreMode, all on the DNA binding domain of the p53 protein (**Supplementary Figure 10a**). Among those regions, sites 291 and 292 are essential post-translational modification sites for p53 ubiquitination and subsequent degradation^70^. A previous study showed the variants at sites 121 and 290-292 increased the ability to induce apoptosis in cultured cells^71^ (**Supplementary Figure 10a**).

PreMode also identified several GoF enriched regions in *RET* and *KCNJ11*. In *RET*, PreMode identified GoF enriched regions both in the kinase domain and regulatory signaling domain near the extracellular binding sites (**Supplementary Figure 10b**). In *KCNJ11*, the regions are located at transmembrane domains and cytoplasmic domains (48-54, 158-182, 200-210,225-231, 320-331, PDB: 6C3O) (**Supplementary Figure 10c**). The region spanning positions 160-179 forms the core part of potassium channel in the tetramer (**Supplementary Figure 11**), especially residues L164 and F168 that form the inner helix gate^72^, and the regions spanning position 48-54, 179-182, 328-331 forms the ATP/ADP binding pocket (**Supplementary Figure 11**).

*PTEN* is a tumor suppressor gene and an essential gene in fetal development. Loss of function in *PTEN* can lead to multiple syndromes. As shown by deep mutational scan experiments^33,73^, some variants in *PTEN* can have different impacts on stability and enzyme activities. We applied PreMode on 80% of the data from both functional assays to predict the effect of all possible variants in *PTEN*. Although most of variants decrease both the protein stability and enzyme function, PreMode identified variants that only disrupt stability but not enzyme function (**Supplementary Figure 12a**, blue), and such variants are located all over the protein. Similarly, PreMode identified variants that only disrupt enzyme function but not stability, all located in the phosphatase domain (**Supplementary Figure 12a**, red). These variants may have dominant negative effects. In fact, three known dominant negative variants in *PTEN* (C124S, G129E and R130G)^73,74^ were successfully predicted by PreMode to maintain stability while causing loss of enzyme function, among which G129E was not in the training data. PreMode identified 4 regions enriched for such variants (**Supplementary Figure 12a**, pink). Further analysis showed those regions are spatially close to each other and form the enzyme pocket around the phosphatase site (**Supplementary Figure 12b**). Notably, PreMode can identify similar dominant-negative variants enriched in a region with only 20% (398 points) of the dataset (**Supplementary Figure 13**).

### PreMode trained with active learning allows efficient few-shot transfer

Deep mutational scan and directed evolution-based protein design experiments often incur expensive time costs upon scaling up. We asked if PreMode can be used to optimize experimental design by active learning^75^. In an active learning framework, PreMode was iteratively trained on a set of experimental data and prioritized the rest of the variants with largest predicted variance to be measured in the next rounds of experiments (**Methods**). We applied this framework to the protein design of green fluorescence protein (GFP) on fluorescence strength. PreMode was able to predict the fitness landscape of GFP with spearman correlation above state-of-the-art performance (0.69)^76–78^ to the experimental data using only 40% of the training data by adaptive learning, which is much more efficient than randomly subsampling data (**Supplementary Figure 14**)

## Discussion

Previous methods for predicting the effect of missense variant have been focused on pathogenicity, which is a binary label. However, different pathogenic variants in the same gene can have different modes of action, i.e., change the protein function in different or sometimes opposite ways. It is challenging to predict mode-of-action because it is gene specific that varies across genes depending on the functions of encoded proteins, yet there is very limited amount of data in individual genes. In this study, we addressed this issue with a new deep learning method, PreMode, that enables pretraining on large pathogenic datasets across genes and then transfer learning in specific genes that have small number of variants with known mode-of-action.

To generate the training data for PreMode as well as understand the biochemical and evolutionary differences of gain/loss-of-function variants, we curated and characterized the largest-to-date missense variants that are known to act through different modes. Gain of function variants tend to be located in low pLDDT regions and surfaces in Alphafold2 predicted structures. Those regions are likely to be intrinsically disordered^79^ which implies conformational heterogeneity and dynamics that could be associated with protein binding and regulatory activities. On the contrary, loss of function variants tend to increase protein folding energy, suggesting that some of these variants destabilize protein structures, while gain-of-function variants tend to preserve protein folding integrity^35^. Overall, these findings showed how variant effects are associated with the protein structures and functions qualitatively. Gain-of-function variants often act through preserving overall protein structure while operating through diverse mechanisms by targeting specific structural domains, whereas LoF variants often destabilize protein structures and tend to be distributed across structured regions. However, this overall trend does not apply to some protein families, for examples GoF variants are more enriched in alpha helixes that forms the transmembrane domain in the ion channel proteins. This highlights the need for protein-specific rather than genome-wide Mode-of-Action predictions. Additionally, available data are heavily uneven across genes, making deep learning algorithms easily stuck at local minimum and gene properties to distinguish genes that tend to act through G/LoF mechanisms while hard to distinguish G/LoF variants within same gene.

To predict protein-specific mode-of-action from the sequence and structural context, we designed a deep learning method, PreMode. PreMode encodes the variant effects with implicit representation of biochemical properties and evolutionary information from protein language model embeddings, and explicit representations of protein structures via SE(3)-equivariant neural networks. PreMode performs gene-specific mode-of-action predictions through a genome-wide pretrain stage and a gene-specific transfer learning stage based on the hypothesis that the sequence and structural context that are informative for pathogenicity should also be informative for mode-of-action predictions. We selected 17 genes with sufficient deep mutation scan data or labeled gain/loss-of-function missense variants for model development and evaluation. At the molecular level, PreMode can simultaneously predict the multi-dimensional biochemical impact of single missense variants, which can reveal potential dominant negative variants that reduce protein function but maintain stability. We showed that PreMode is efficient at transfer learning and that it can capture the fitness landscape of all possible variants within one protein using around 40% of mutagenesis data. At genetic level, PreMode can efficiently utilize a small amount labeled data (a few dozens) to accurately distinguish G/LoF variants with an AUC of around 0.80 in most proteins. Additionally, we showed PreMode’s utility in deep mutational scan experiments and protein engineering. First, PreMode can improve efficiency of deep mutational scan experiments by detecting noisy data points in single measurements. Second, PreMode can be applied to unmeasured genes by fine tuning on the stability deep mutational experiments. Finally, PreMode can facilitate mutagenesis-based protein directed evolution through adaptive learning by efficiently lowering library sizes.

PreMode currently predicts a binary label as gain or loss of function. This is a limitation as it does not capture the complexity of protein functional changes. For instance, ion channel proteins undergo complex conformation changes and regulation to perform normal physiological functions^29^. Large scale functional studies of these genes may provide additional data that enable training of improved models. Furthermore, gain/loss of function could be further divided into amorph, hypomorph, hypermorph, antimorph and neomorph according to muller’s morphs^80^. Accordingly, additional labeled data may facilitate the training of more accurate and comprehensive models.

PreMode can be potentially improved by considering protein dynamics, as ablation experiments showed it has lower performance for genes where both G/LoF variants are located in regions with relatively low pLDDT values. Currently PreMode includes features representing a static protein structure, which is not sufficient to model the variant effects in those regions.

## Data and code availability

All the data and code used in model training and analysis could be found at GitHub: https://github.com/ShenLab/PreMode. We also provided all the model weights and all mode-of-action predictions for genes *PTEN, BRAF, KCNJ11, TP53 and RET* in the repository.

## Acknowledgement

This work was supported by NIH grants (R35GM149527, P50HD109879) and Simons Foundation (SFARI #1019623). We thank Dr. Mohammed AlQuraishi, Dr. David Knowles, Dr. Haicang Zhang, Dr. Haiqing Zhao, Xi Fu, Dr. Xiao Fan, Dr. Na Zhu and Dr. Chao Hou for helpful discussions and suggestions.

## Methods

### Training and testing datasets

We curated labeled pathogenic/benign variants, gain/loss of function variants, deep mutational scan experiments from public databases and publications. For gain/loss of function prediction tasks, we collected: 765 gain and 4,571 loss of function missense variants from Barak, et al^44^; 669 gain and 1,232 loss of function variants from Clinical Knowledge Base^52^; 199 gain and 1,506 loss of function variants from cBioportal cancer hotspots^49^; 56 gain and 57 loss of function variants in *ABCC8, GCK, KCNJ11* from Flanagan, et al.^47^; 45 gain and 7 loss of function variants in *STAT1* from Kagawa, et al.^48^; 309 gain and 516 loss of function variants from Heyne, et al^36^. More specifically, for variants in cBioportal, we first calculate cancer hotspots based on existing algorithms^58^, then annotate 199 variants in hotspots of 27 oncogenic genes as gain of function and 1506 variants in 248 tumor suppressor genes as loss of function based on COSMIC database^50^. We excluded genes with multiple cancer roles.

As the gain and loss of function variants were extremely biased across genes, we didn’t split the training and testing dataset in common machine learning manner but split by protein-wise manner with the following steps:

1. For each gene, we select 20% of GoF and 20% of LoF variants as testing. We use the rest variants in the same gene as training. We did the split 5 times.
2. We only kept genes with more than 15 GoF variants and 15 LoF variants for model evaluation. For comparison to LoGoFunc, we split the same amount of testing data that not in LoGoFunc’s data, and use the rest of training. We did the split 5 times.

For comparison to FuNCion, we split the training/testing data based on their split.

For predicting pathogenicity, we collected 148,324 variants for training, including: 51,494 benign variants from PrimateAI^17^ and 32,350 non-overlapping benign variants from ClinVar with at least one-star non-conflict submits that labeled as “benign” or “likely-benign”; 64,480 pathogenic variants from ClinVar database with at least one-star non-conflict submits^7^, non-overlapping variants from HGMD^40^. We collected 1,066 variants for testing, including 533 pathogenic missense variants from somatic missense hotspots in 153 cancer driver genes that not annotated above; and 533 benign variants from the same genes randomly selected from ClinVar and PrimateAI not overlapped with training dataset.

For deep mutational scan assays, we collected datasets of *PTEN, SNCA, CCR5, CXCR4, NUDT15, CYP2C9, GCK, ASPA* from MAVEDB^6^ (**Supplementary Table 2**).

### Input features

For a missense variant of interest, PreMode considers a 250 amino acid window flanking the variant position and the residues as nodes, and constructs two graphs based on its protein context; the first graph *G*_1_ is a non-directed star graph that connect only the variant node and the other nodes. The second graph *G*_2_ is a non-directed K-nearest neighbor graph that connect each node with its neighbors based on 3D Euclidean distance of the alpha carbon atoms.

Each node has a set of invariant features and structure features. The invariant features include: Embeddings from the last layer of ESM2 (650M)^25^ (d=1280); Dssp^53,54^ annotated secondary structure and torsion angles from the AlphaFold2 predicted protein structures (d=12); pLDDT^24^ for the AlphaFold2 prediction; Amino acid frequencies from MSA of 199 species that reflects evolutionary conservation (d=20). The variant node has additional invariant feature of the embedding of alternate amino acid. The structure features include a set of Euclidean vectors from the alpha carbon to all other non-hydrogen atoms of side chain (d=3×35). If an atom does not exist in the side chain, then it is set as 0.

Each edge has a set of invariant features and structure features. The invariant features include, weighted covariance matrix between the amino acid frequencies of two residue sites in multiple sequence alignments (MSA) of 199 species^19^ (ALGORITHM 1); Euclidean distances of beta carbons between two residues (For glycine we use alpha carbon), transformed by exponential smearing functions (ALGORITHM 2); The contact strength predicted by MSA transformer. The structure features include the Euclidean vector of beta carbons between two residues.

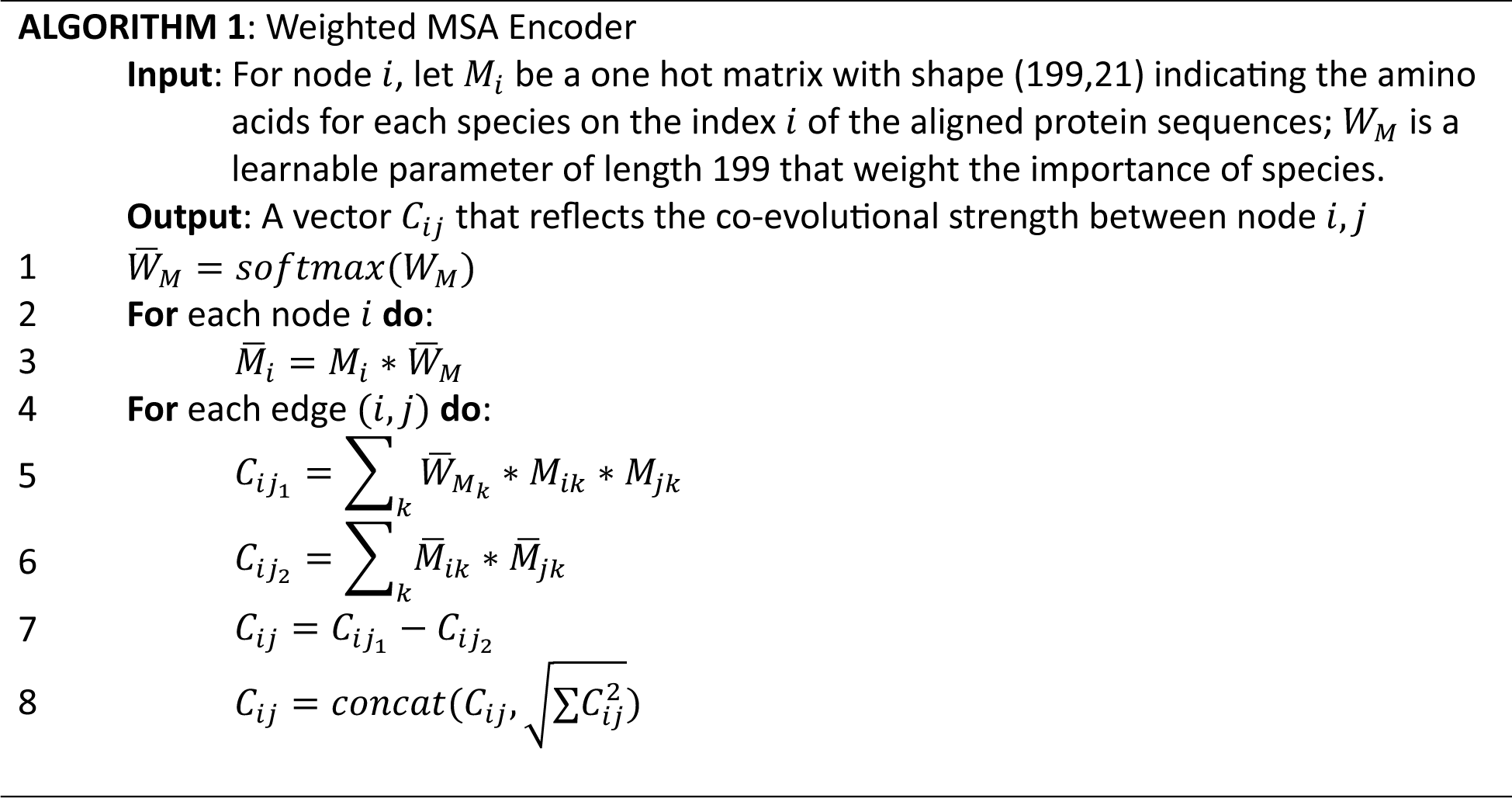

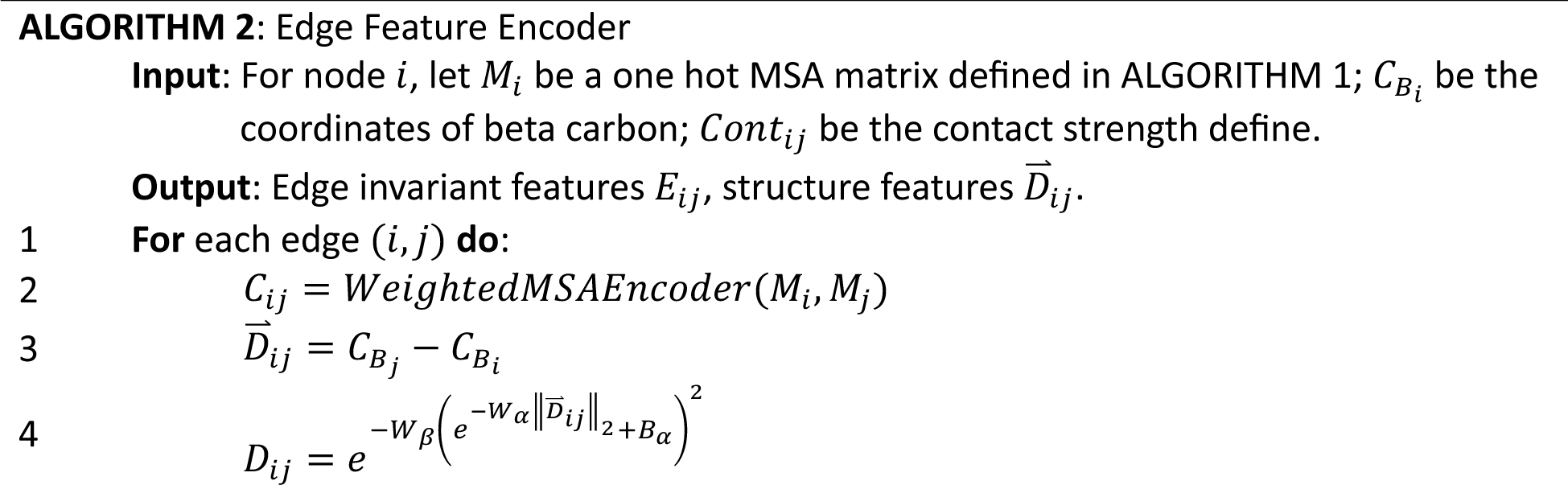
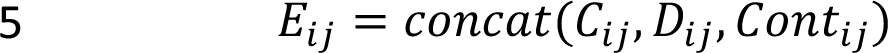

### Model architectures

PreMode is an SE(3)-equivariant graph attention neural network with 4 layers. First layer is a feature embedding layer that encodes the input features into latent dimensions. This layer was designed separately for invariant features and structure features. For invariant features, PreMode uses GeLU^81^ as the activation function and linear layer of 512 dimensions with weights *W*_i_ and bias *B*_i_ as output. The variant node has an additional embedding layer with weights *B*_*i*_ and bias *B*_*i*_ to incorporate with the alternate amino acid embeddings (ALGORITHM 3). For structure features, PreMode uses a linear layer with weights *W*_*s*_ but without bias term to ensure the equivariance. The latent dimension is 32 (**Supplementary Figure 1**). For edge features, PreMode encodes the pairwise MSA, Euclidean distances, contact predictions, relative positions into a 444-dim invariant feature vector.

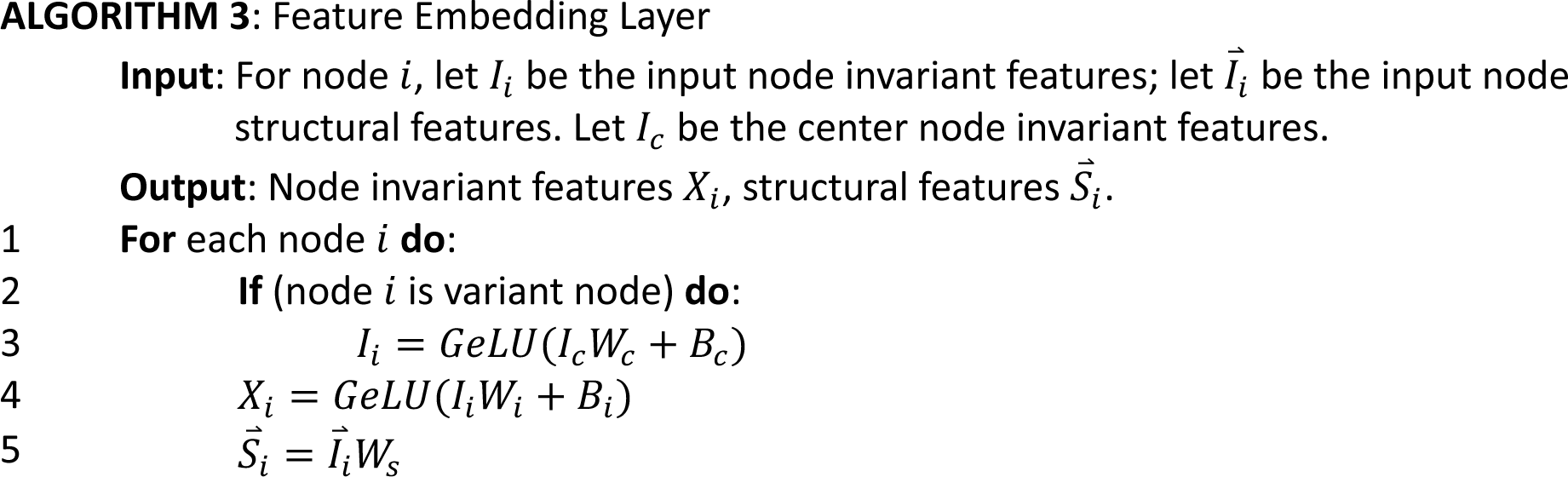

Second and third layers are equivariant graph attention layers. We calculate the attention values only between the connected nodes in star or KNN graphs defined in the features section. We used a modified SE(3)-equivariant attention mechanism inspired from torchmd-net^82^ and gMVP^19^ to calculate the attention values that takes co-evolutional evolution, structure features into consideration. The invariant and structure features were updated separately but share information across each other based on the attention values to maintain equivariance (ALGORITHM 4). Those two layers were designed to capture the biochemical context for the residue of interest, where the second layer focuses more on the first order impact of amino acid change to all other residues, and the third layer implicitly models the second order consequences.

The last layer is a graph attention layer that designed to summarize the overall impact of the variant to protein. It only takes the invariant embeddings output from the third layer to calculate the attention values between center node to other nodes while don’t take structural features.

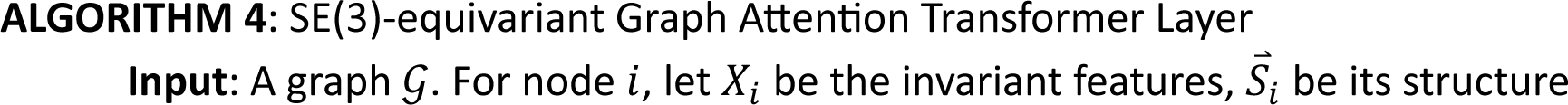
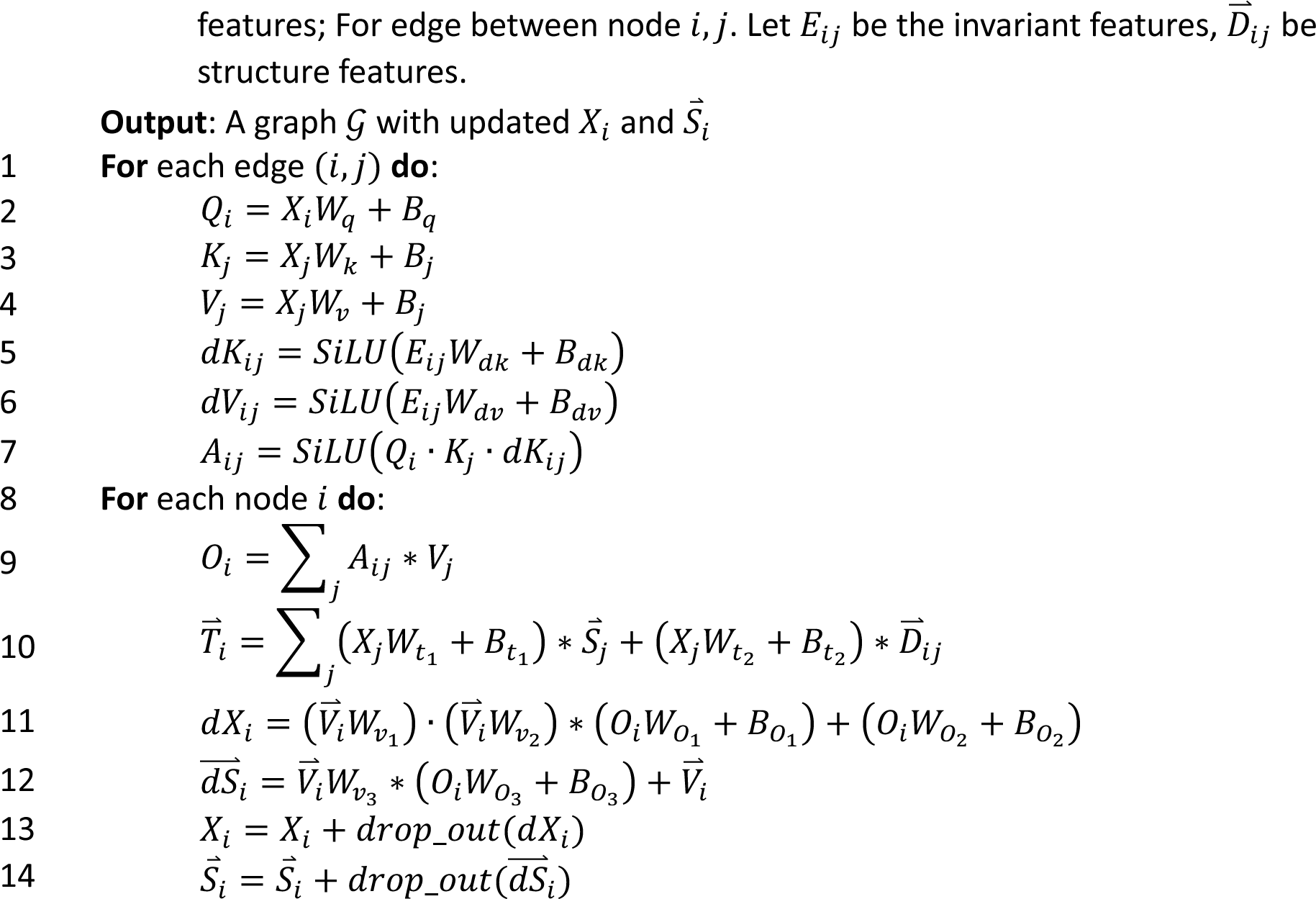

### Model training and testing

We trained PreMode with Adam^83^ algorithm. For pretrain on pathogenicity dataset, we set learning rate to 1e-4, batch size to 256 and trained 20 epochs. We randomly selected 5% of training data as validation and calculated the loss on it every 250 steps. We trained the model under 500 steps of warmup stage, with linear learning rate increase until maximum learning rate 1e-4, followed by the learning rate decay stage, where we decreased the learning rate by 0.8 if observed a plateau on the validation loss until the minimum learning rate 1e-6. After training, we selected the model with minimum validation loss for testing. The training process took about 50h under 4 NVIDIA A40 GPUs.

For transfer learning on molecular mode-of-action tasks, we set the batch size to 8, training epochs to 40. We trained the model with 200 steps of warmup stage steps, evaluated validation loss every 400 steps and set the minimum learning rate to 1e-7 in the learning rate decay stage. After training, we selected the model with minimum validation loss for testing. The training process took about 4∼6h under 1 NVIDIA A40 GPU depending on the dataset size.

For transfer learning on genetic mode-of-action tasks, we set everything same as the molecular mode-of-action tasks, except that we trained the model with 20 epochs and evaluate validation loss every 5 steps in the learning rate decay stage. For validation dataset, we randomly split training data into 4 parts and did 4-fold cross-validation. After training, we selected the models with minimum validation loss under each cross-validation and ensemble them by averaging their predictions. We trained two ensemble models with window size of 251 and 1251 under this same scheme, and selected the ensemble model with best overall AUC on whole training data for evaluation on testing data. The training process took about 20min∼1h under 1 NVIDIA A40 GPU depending on the dataset size.

For pretrain and transfer learning on genetic mode-of-action tasks, we calculated the AUROC (area under recall receiver operating characteristic curve) on testing data to assess the performances of models. For pretrain task, we only calculated one AUROC value. For transfer learning tasks, we calculated AUROC for all five random training/testing splits and reported the mean and standard error. For molecular mode-of-action tasks, we calculated the spearman correlation between model prediction and experimental measurement on testing data. We calculated spearman correlation for all five random training/testing splits and reported the mean and standard error.

### Baseline Methods and ablation analysis

For both molecular and genetics level mode-of-action predictions, we created several baseline methods to compare PreMode with. First, we built a single layer perceptron (SLP) model on top of the ESM embeddings (ESM + SLP). This model took all invariant features (ESM2 embeddings, amino acid changes, conservation in MSA) that same as PreMode as inputs, followed by one linear layer and GeLU activation layer. Next, we implemented several ablation analyses on PreMode. For structure feature ablation, we replaced all structural features with zeros, and constructed the star and KNN graphs based on the 1D-distance on the sequences. For ESM embedding feature ablation, we replaced the ESM embeddings with one-hot encodings of 20 amino acids. For MSA feature ablation, we removed all MSA inputs and the MSA attention module. All the models above were trained under same training configurations as PreMode at both pretrain and transfer learning stages. Lastly, we implemented low-capacity machine learning models using only biochemical properties as inputs, including conservation, secondary structure, reference, and alternate amino acid identities, pLDDT and ddG. For molecular-level mode-of-action predictions, we implemented it with elastic net linear regression model while for genetics-level mode-of-action predictions, we implemented it with random forest classification model. To compare the overall performances of models, for molecular mode-of-action predictions, we weighted average the spearman correlation for each gene based on their dataset sizes. For genetics mode-of-action predictions, we first calculate the effective data set sizes as the harmonic average of gain and loss of function variants in each gene, then weighted average the AUC across genes.

### Curation of prediction scores from other methods

In the pathogenicity prediction comparison, we directly obtained the prediction scores for PrimateAI, EVE, REVEL from dbNSFP (v4.4a)^84^. For gMVP, AlphaMissense, ESM1b, we obtained the prediction scores from their original publications^19,20,28^.

In the molecular mode-of-action prediction comparison, we selected three models with top performances reported in Hsu, et al.^59^, Augmented ESM1b, Augmented EVmutation and Augmented Unirep. We didn’t compare with Augmented DeepSequence due to errors in the publicly available codes. Instead, we trained the augmented model using evolutionary density score from EVE^16^, as both models were based on variational autoencoders on MSA. We trained and tested those models using the same MSA and protein sequence inputs as well as same training and testing data as PreMode.

In the genetic mode-of-action prediction comparison, we obtained LoGoFunc prediction scores from https://itanlab.shinyapps.io/goflof/. For FuNCion, we obtained their codes and training/testing split information from their GitHub page (https://github.com/heyhen/funNCion). We compared PreMode and their model’s performance using their training/testing split as well as ours. For FuNCion, their original implementation of gradient boosting machine learning method in R “caret” package will raise errors under small sample sizes, we reimplemented the gradient boosting method in python “sklearn” package and reported both AUCs.

### Subsample of datasets

For deep mutational scan datasets, we subsampled the data to investigate how many points were required for sufficient adaptation to the tasks in transfer learning. For each of the multi-dimensional assays, we randomly subsampled training data to 10%, 20%, 40%, 60%, 80% of whole datasets 5 times with different random seeds and testing on the same 20% of whole datasets. We performed PreMode on each of the training data and evaluate the performance.

### In silico saturated mutagenesis experiments

We did in silico saturated mutagenesis experiments for *BRAF*, *TP53*, *KCNJ11*, *RET*, and *PTEN*. For each gene, we calculate two predictions *r* (pathogenicity score) and θ (Gain/Loss of function score) with pretrained model and corresponding transfer-learning models, respectively. For θ, there are 5 predicted scores from models trained on 5 training/testing splits under different random seeds. We selected the models with minimum validation loss under each training/testing split and ensembled the predictions by voting with a simple random forest model.

### Active Learning experiments

We split the deep mutational scan experiment datasets to ∼40% of training data, ∼10% of validation data and ∼50% of testing data the same way as previous methods^76–78^. Within the training data, we first performed PreMode transfer learning on 10% of the randomly selected data, then evaluated PreMode on the rest of 90% of training data. PreMode will output both predicted values as well as model confidence values. Then we selected the top 10% of data among the rest of training data where PreMode was most unconfident and added them to the next round of training. We repeated this procedure until all the training data were used.

## References

1. Homsy, J., Zaidi, S., Shen, Y., Ware, J.S., Samocha, K.E., Karczewski, K.J., DePalma, S.R., McKean, D., Wakimoto, H., Gorham, J., et al. (2015). De novo mutations in congenital heart disease with neurodevelopmental and other congenital anomalies. Science 350, 1262–1266. 10.1126/science.aac9396.

2. Jin, S.C., Homsy, J., Zaidi, S., Lu, Q., Morton, S., DePalma, S.R., Zeng, X., Qi, H., Chang, W., Sierant, M.C., et al. (2017). Contribution of rare inherited and de novo variants in 2,871 congenital heart disease probands. Nat Genet 49, 1593–1601. 10.1038/ng.3970.

3. Huang, K.L., Mashl, R.J., Wu, Y., Ritter, D.I., Wang, J., Oh, C., Paczkowska, M., Reynolds, S., Wyczalkowski, M.A., Oak, N., et al. (2018). Pathogenic Germline Variants in 10,389 Adult Cancers. Cell 173, 355–370 e314. 10.1016/j.cell.2018.03.039.

4. Satterstrom, F.K., Kosmicki, J.A., Wang, J., Breen, M.S., De Rubeis, S., An, J.Y., Peng, M., Collins, R., Grove, J., Klei, L., et al. (2020). Large-Scale Exome Sequencing Study Implicates Both Developmental and Functional Changes in the Neurobiology of Autism. Cell 180, 568–584 e523. 10.1016/j.cell.2019.12.036.

5. Zhou, X., Feliciano, P., Shu, C., Wang, T., Astrovskaya, I., Hall, J.B., Obiajulu, J.U., Wright, J.R., Murali, S.C., Xu, S.X., et al. (2022). Integrating de novo and inherited variants in 42,607 autism cases identifies mutations in new moderate-risk genes. Nat Genet 54, 1305–1319. 10.1038/s41588-022-01148-2.

6. Esposito, D., Weile, J., Shendure, J., Starita, L.M., Papenfuss, A.T., Roth, F.P., Fowler, D.M., and Rubin, A.F. (2019). MaveDB: an open-source platform to distribute and interpret data from multiplexed assays of variant effect. Genome Biol 20, 223. 10.1186/s13059-019-1845-6.

7. Landrum, M.J., Lee, J.M., Benson, M., Brown, G.R., Chao, C., Chitipiralla, S., Gu, B., Hart, J., Hoffman, D., Jang, W., et al. (2018). ClinVar: improving access to variant interpretations and supporting evidence. Nucleic Acids Res 46, D1062–D1067. 10.1093/nar/gkx1153.

8. Romero, P.A., and Arnold, F.H. (2009). Exploring protein fitness landscapes by directed evolution. Nat Rev Mol Cell Biol 10, 866–876. 10.1038/nrm2805.

9. Yang, K.K., Wu, Z., and Arnold, F.H. (2019). Machine-learning-guided directed evolution for protein engineering. Nat Methods 16, 687–694. 10.1038/s41592-019-0496-6.

10. Wang, Y., Xue, P., Cao, M., Yu, T., Lane, S.T., and Zhao, H. (2021). Directed Evolution: Methodologies and Applications. Chem Rev 121, 12384–12444. 10.1021/acs.chemrev.1c00260.

11. Adzhubei, I.A., Schmidt, S., Peshkin, L., Ramensky, V.E., Gerasimova, A., Bork, P., Kondrashov, A.S., and Sunyaev, S.R. (2010). A method and server for predicting damaging missense mutations. Nature methods 7, 248–249. 10.1038/nmeth0410-248.

12. Ng, P.C., and Henikoff, S. (2003). SIFT: Predicting amino acid changes that affect protein function. Nucleic Acids Res 31, 3812–3814. 10.1093/nar/gkg509.

13. Kircher, M., Witten, D.M., Jain, P., O’Roak, B.J., Cooper, G.M., and Shendure, J. (2014). A general framework for estimating the relative pathogenicity of human genetic variants. Nat Genet 46, 310–315. 10.1038/ng.2892.

14. Ioannidis, N.M., Rothstein, J.H., Pejaver, V., Middha, S., McDonnell, S.K., Baheti, S., Musolf, A., Li, Q., Holzinger, E., Karyadi, D., et al. (2016). REVEL: An Ensemble Method for Predicting the Pathogenicity of Rare Missense Variants. Am J Hum Genet 99, 877–885. 10.1016/j.ajhg.2016.08.016.

15. Samocha, K.E., Kosmicki, J.A., Karczewski, K.J., O’Donnell-Luria, A.H., Pierce-Hoffman, E., MacArthur, D.G., Neale, B.M., and Daly, M.J. (2017). Regional missense constraint improves variant deleteriousness prediction. bioRxiv, 148353. 10.1101/148353.

16. Frazer, J., Notin, P., Dias, M., Gomez, A., Min, J.K., Brock, K., Gal, Y., and Marks, D.S. (2021). Disease variant prediction with deep generative models of evolutionary data. Nature 599, 91–95. 10.1038/s41586-021-04043-8.

17. Sundaram, L., Gao, H., Padigepati, S.R., McRae, J.F., Li, Y., Kosmicki, J.A., Fritzilas, N., Hakenberg, J., Dutta, A., Shon, J., et al. (2018). Predicting the clinical impact of human mutation with deep neural networks. Nat Genet 50, 1161–1170. 10.1038/s41588-018-0167-z.

18. Gao, H., Hamp, T., Ede, J., Schraiber, J.G., McRae, J., Singer-Berk, M., Yang, Y., Dietrich, A.S.D., Fiziev, P.P., Kuderna, L.F.K., et al. (2023). The landscape of tolerated genetic variation in humans and primates. Science 380, eabn8153. 10.1126/science.abn8197.

19. Zhang, H., Xu, M.S., Fan, X., Chung, W.K., and Shen, Y. (2022). Predicting functional effect of missense variants using graph attention neural networks. Nature Machine Intelligence 4, 1017–1028. 10.1038/s42256-022-00561-w.

20. Cheng, J., Novati, G., Pan, J., Bycroft, C., Zemgulyte, A., Applebaum, T., Pritzel, A., Wong, L.H., Zielinski, M., Sargeant, T., et al. (2023). Accurate proteome-wide missense variant effect prediction with AlphaMissense. Science 381, eadg7492. 10.1126/science.adg7492.

21. Feng, B.J. (2017). PERCH: A Unified Framework for Disease Gene Prioritization. Hum Mutat 38, 243–251. 10.1002/humu.23158.

22. Jagadeesh, K.A., Wenger, A.M., Berger, M.J., Guturu, H., Stenson, P.D., Cooper, D.N., Bernstein, J.A., and Bejerano, G. (2016). M-CAP eliminates a majority of variants of uncertain significance in clinical exomes at high sensitivity. Nat Genet 48, 1581–1586. 10.1038/ng.3703.

23. UniProt, C. (2021). UniProt: the universal protein knowledgebase in 2021. Nucleic Acids Res 49, D480–D489. 10.1093/nar/gkaa1100.

24. Jumper, J., Evans, R., Pritzel, A., Green, T., Figurnov, M., Ronneberger, O., Tunyasuvunakool, K., Bates, R., Zidek, A., Potapenko, A., et al. (2021). Highly accurate protein structure prediction with AlphaFold. Nature 596, 583–589. 10.1038/s41586-021-03819-2.

25. Lin, Z., Akin, H., Rao, R., Hie, B., Zhu, Z., Lu, W., Smetanin, N., Verkuil, R., Kabeli, O., Shmueli, Y., et al. (2023). Evolutionary-scale prediction of atomic-level protein structure with a language model. Science 379, 1123–1130. 10.1126/science.ade2574.

26. Brandes, N., Ofer, D., Peleg, Y., Rappoport, N., and Linial, M. (2022). ProteinBERT: a universal deep-learning model of protein sequence and function. Bioinformatics 38, 2102–2110. 10.1093/bioinformatics/btac020.

27. Meier, J., Rao, R., Verkuil, R., Liu, J., Sercu, T., and Rives, A. (2021). Language models enable zero-shot prediction of the effects of mutations on protein function. bioRxiv, 2021.2007.2009.450648. 10.1101/2021.07.09.450648.

28. Brandes, N., Goldman, G., Wang, C.H., Ye, C.J., and Ntranos, V. (2023). Genome-wide prediction of disease variant effects with a deep protein language model. Nat Genet 55, 1512–1522. 10.1038/s41588-023-01465-0.

29. Sanders, S.J., Campbell, A.J., Cottrell, J.R., Moller, R.S., Wagner, F.F., Auldridge, A.L., Bernier, R.A., Catterall, W.A., Chung, W.K., Empfield, J.R., et al. (2018). Progress in Understanding and Treating SCN2A-Mediated Disorders. Trends Neurosci 41, 442–456. 10.1016/j.tins.2018.03.011.

30. Wolff, M., Brunklaus, A., and Zuberi, S.M. (2019). Phenotypic spectrum and genetics of SCN2A-related disorders, treatment options, and outcomes in epilepsy and beyond. Epilepsia 60 Suppl 3, S59–S67. 10.1111/epi.14935.

31. Matreyek, K.A., Starita, L.M., Stephany, J.J., Martin, B., Chiasson, M.A., Gray, V.E., Kircher, M., Khechaduri, A., Dines, J.N., Hause, R.J., et al. (2018). Multiplex assessment of protein variant abundance by massively parallel sequencing. Nat Genet 50, 874–882. 10.1038/s41588-018-0122-z.

32. Suiter, C.C., Moriyama, T., Matreyek, K.A., Yang, W., Scaletti, E.R., Nishii, R., Yang, W., Hoshitsuki, K., Singh, M., Trehan, A., et al. (2020). Massively parallel variant characterization identifies NUDT15 alleles associated with thiopurine toxicity. Proc Natl Acad Sci U S A 117, 5394–5401. 10.1073/pnas.1915680117.

33. Mighell, T.L., Evans-Dutson, S., and O’Roak, B.J. (2018). A Saturation Mutagenesis Approach to Understanding PTEN Lipid Phosphatase Activity and Genotype-Phenotype Relationships. Am J Hum Genet 102, 943–955. 10.1016/j.ajhg.2018.03.018.

34. Lage, K. (2014). Protein-protein interactions and genetic diseases: The interactome. Biochim Biophys Acta 1842, 1971–1980. 10.1016/j.bbadis.2014.05.028.

35. Gerasimavicius, L., Livesey, B.J., and Marsh, J.A. (2022). Loss-of-function, gain-of-function and dominant-negative mutations have profoundly different effects on protein structure. Nat Commun 13, 3895. 10.1038/s41467-022-31686-6.

36. Heyne, H.O., Baez-Nieto, D., Iqbal, S., Palmer, D.S., Brunklaus, A., May, P., Epi, C., Johannesen, K.M., Lauxmann, S., Lemke, J.R., et al. (2020). Predicting functional effects of missense variants in voltage-gated sodium and calcium channels. Science translational medicine 12. 10.1126/scitranslmed.aay6848.

37. Lester, H.A., and Karschin, A. (2000). Gain of function mutants: ion channels and G protein-coupled receptors. Annu Rev Neurosci 23, 89–125. 10.1146/annurev.neuro.23.1.89.

38. Li, Y., Zhang, Y., Li, X., Yi, S., and Xu, J. (2019). Gain-of-Function Mutations: An Emerging Advantage for Cancer Biology. Trends Biochem Sci 44, 659–674. 10.1016/j.tibs.2019.03.009.

39. Wang, L.H., Wu, C.F., Rajasekaran, N., and Shin, Y.K. (2018). Loss of Tumor Suppressor Gene Function in Human Cancer: An Overview. Cell Physiol Biochem 51, 2647–2693. 10.1159/000495956.

40. Stenson, P.D., Mort, M., Ball, E.V., Evans, K., Hayden, M., Heywood, S., Hussain, M., Phillips, A.D., and Cooper, D.N. (2017). The Human Gene Mutation Database: towards a comprehensive repository of inherited mutation data for medical research, genetic diagnosis and next-generation sequencing studies. Hum Genet 136, 665–677. 10.1007/s00439-017-1779-6.

41. Swan, H., Amarouch, M.Y., Leinonen, J., Marjamaa, A., Kucera, J.P., Laitinen-Forsblom, P.J., Lahtinen, A.M., Palotie, A., Kontula, K., Toivonen, L., et al. (2014). Gain-of-function mutation of the SCN5A gene causes exercise-induced polymorphic ventricular arrhythmias. Circ Cardiovasc Genet 7, 771–781. 10.1161/CIRCGENETICS.114.000703.

42. Blanchard, M.G., Willemsen, M.H., Walker, J.B., Dib-Hajj, S.D., Waxman, S.G., Jongmans, M.C., Kleefstra, T., van de Warrenburg, B.P., Praamstra, P., Nicolai, J., et al. (2015). De novo gain-of-function and loss-of-function mutations of SCN8A in patients with intellectual disabilities and epilepsy. J Med Genet 52, 330–337. 10.1136/jmedgenet-2014-102813.

43. McDonell, L.M., Kernohan, K.D., Boycott, K.M., and Sawyer, S.L. (2015). Receptor tyrosine kinase mutations in developmental syndromes and cancer: two sides of the same coin. Hum Mol Genet 24, R60–66. 10.1093/hmg/ddv254.

44. Sevim Bayrak, C., Stein, D., Jain, A., Chaudhary, K., Nadkarni, G.N., Van Vleck, T.T., Puel, A., Boisson-Dupuis, S., Okada, S., Stenson, P.D., et al. (2021). Identification of discriminative gene-level and protein-level features associated with pathogenic gain-of-function and loss-of-function variants. Am J Hum Genet 108, 2301–2318. 10.1016/j.ajhg.2021.10.007.

45. Zhang, C., Liu, J., Xu, D., Zhang, T., Hu, W., and Feng, Z. (2020). Gain-of-function mutant p53 in cancer progression and therapy. J Mol Cell Biol 12, 674–687. 10.1093/jmcb/mjaa040.

46. Stein, D., Kars, M.E., Wu, Y., Bayrak, C.S., Stenson, P.D., Cooper, D.N., Schlessinger, A., and Itan, Y. (2023). Genome-wide prediction of pathogenic gain-and loss-of-function variants from ensemble learning of a diverse feature set. Genome Med 15, 103. 10.1186/s13073-023-01261-9.

47. Flanagan, S.E., Patch, A.M., and Ellard, S. (2010). Using SIFT and PolyPhen to predict loss-of-function and gain-of-function mutations. Genet Test Mol Biomarkers 14, 533–537. 10.1089/gtmb.2010.0036.

48. Kagawa, R., Fujiki, R., Tsumura, M., Sakata, S., Nishimura, S., Itan, Y., Kong, X.F., Kato, Z., Ohnishi, H., Hirata, O., et al. (2017). Alanine-scanning mutagenesis of human signal transducer and activator of transcription 1 to estimate loss-or gain-of-function variants. J Allergy Clin Immunol 140, 232–241. 10.1016/j.jaci.2016.09.035.

49. Gao, J., Aksoy, B.A., Dogrusoz, U., Dresdner, G., Gross, B., Sumer, S.O., Sun, Y., Jacobsen, A., Sinha, R., Larsson, E., et al. (2013). Integrative analysis of complex cancer genomics and clinical profiles using the cBioPortal. Sci Signal 6, pl1. 10.1126/scisignal.2004088.

50. Tate, J.G., Bamford, S., Jubb, H.C., Sondka, Z., Beare, D.M., Bindal, N., Boutselakis, H., Cole, C.G., Creatore, C., Dawson, E., et al. (2019). COSMIC: the Catalogue Of Somatic Mutations In Cancer. Nucleic Acids Res 47, D941–D947. 10.1093/nar/gky1015.

51. Chang, M.T., Bhattarai, T.S., Schram, A.M., Bielski, C.M., Donoghue, M.T.A., Jonsson, P., Chakravarty, D., Phillips, S., Kandoth, C., Penson, A., et al. (2018). Accelerating Discovery of Functional Mutant Alleles in Cancer. Cancer Discov 8, 174–183. 10.1158/2159-8290.CD-17-0321.

52. Patterson, S.E., Statz, C.M., Yin, T., and Mockus, S.M. (2019). Utility of the JAX Clinical Knowledgebase in capture and assessment of complex genomic cancer data. NPJ Precis Oncol 3, 2. 10.1038/s41698-018-0073-y.

53. Joosten, R.P., te Beek, T.A., Krieger, E., Hekkelman, M.L., Hooft, R.W., Schneider, R., Sander, C., and Vriend, G. (2011). A series of PDB related databases for everyday needs. Nucleic Acids Res 39, D411–419. 10.1093/nar/gkq1105.

54. Kabsch, W., and Sander, C. (1983). Dictionary of protein secondary structure: pattern recognition of hydrogen-bonded and geometrical features. Biopolymers 22, 2577–2637. 10.1002/bip.360221211.

55. Valdar, W.S. (2002). Scoring residue conservation. Proteins 48, 227–241. 10.1002/prot.10146.

56. Rao, R.M., Liu, J., Verkuil, R., Meier, J., Canny, J., Abbeel, P., Sercu, T., and Rives, A. (2021). MSA Transformer. In M. Marina, and Z. Tong, eds. Proceedings of the 38th International Conference on Machine Learning. PMLR.

57. Fung, A., Koehl, A., Jagota, M., and Song, Y.S. (2022). The Impact of Protein Dynamics on Residue-Residue Coevolution and Contact Prediction. bioRxiv, 2022.2010.2016.512436. 10.1101/2022.10.16.512436.

58. Chang, M.T., Asthana, S., Gao, S.P., Lee, B.H., Chapman, J.S., Kandoth, C., Gao, J., Socci, N.D., Solit, D.B., Olshen, A.B., et al. (2016). Identifying recurrent mutations in cancer reveals widespread lineage diversity and mutational specificity. Nat Biotechnol 34, 155–163. 10.1038/nbt.3391.

59. Hsu, C., Nisonoff, H., Fannjiang, C., and Listgarten, J. (2022). Learning protein fitness models from evolutionary and assay-labeled data. Nat Biotechnol 40, 1114–1122. 10.1038/s41587-021-01146-5.

60. Hornbeck, P.V., Zhang, B., Murray, B., Kornhauser, J.M., Latham, V., and Skrzypek, E. (2015). PhosphoSitePlus, 2014: mutations, PTMs and recalibrations. Nucleic Acids Res 43, D512-520. 10.1093/nar/gku1267.

61. Wan, P.T., Garnett, M.J., Roe, S.M., Lee, S., Niculescu-Duvaz, D., Good, V.M., Jones, C.M., Marshall, C.J., Springer, C.J., Barford, D., et al. (2004). Mechanism of activation of the RAF-ERK signaling pathway by oncogenic mutations of B-RAF. Cell 116, 855–867. 10.1016/s0092-8674(04)00215-6.

62. Ng, P.K., Li, J., Jeong, K.J., Shao, S., Chen, H., Tsang, Y.H., Sengupta, S., Wang, Z., Bhavana, V.H., Tran, R., et al. (2018). Systematic Functional Annotation of Somatic Mutations in Cancer. Cancer Cell 33, 450–462 e410. 10.1016/j.ccell.2018.01.021.

63. Martinez Fiesco, J.A., Durrant, D.E., Morrison, D.K., and Zhang, P. (2022). Structural insights into the BRAF monomer-to-dimer transition mediated by RAS binding. Nat Commun 13, 486. 10.1038/s41467-022-28084-3.

64. Tran, T.H., Chan, A.H., Young, L.C., Bindu, L., Neale, C., Messing, S., Dharmaiah, S., Taylor, T., Denson, J.P., Esposito, D., et al. (2021). KRAS interaction with RAF1 RAS-binding domain and cysteine-rich domain provides insights into RAS-mediated RAF activation. Nat Commun 12, 1176. 10.1038/s41467-021-21422-x.

65. Mirdita, M., Schutze, K., Moriwaki, Y., Heo, L., Ovchinnikov, S., and Steinegger, M. (2022). ColabFold: making protein folding accessible to all. Nat Methods 19, 679–682. 10.1038/s41592-022-01488-1.

66. Schymkowitz, J., Borg, J., Stricher, F., Nys, R., Rousseau, F., and Serrano, L. (2005). The FoldX web server: an online force field. Nucleic Acids Res 33, W382–388. 10.1093/nar/gki387.

67. Lu, H., Villafane, N., Dogruluk, T., Grzeskowiak, C.L., Kong, K., Tsang, Y.H., Zagorodna, O., Pantazi, A., Yang, L., Neill, N.J., et al. (2017). Engineering and Functional Characterization of Fusion Genes Identifies Novel Oncogenic Drivers of Cancer. Cancer Res 77, 3502–3512. 10.1158/0008-5472.CAN-16-2745.

68. Kiel, C., Benisty, H., Llorens-Rico, V., and Serrano, L. (2016). The yin-yang of kinase activation and unfolding explains the peculiarity of Val600 in the activation segment of BRAF. Elife 5, e12814. 10.7554/eLife.12814.

69. Maloney, R.C., Zhang, M., Jang, H., and Nussinov, R. (2021). The mechanism of activation of monomeric B-Raf V600E. Comput Struct Biotechnol J 19, 3349–3363. 10.1016/j.csbj.2021.06.007.

70. Lee, E.W., Lee, M.S., Camus, S., Ghim, J., Yang, M.R., Oh, W., Ha, N.C., Lane, D.P., and Song, J. (2009). Differential regulation of p53 and p21 by MKRN1 E3 ligase controls cell cycle arrest and apoptosis. EMBO J 28, 2100–2113. 10.1038/emboj.2009.164.

71. Kakudo, Y., Shibata, H., Otsuka, K., Kato, S., and Ishioka, C. (2005). Lack of correlation between p53-dependent transcriptional activity and the ability to induce apoptosis among 179 mutant p53s. Cancer Res 65, 2108–2114. 10.1158/0008-5472.CAN-04-2935.

72. Zhao, C., and MacKinnon, R. (2021). Molecular structure of an open human K(ATP) channel. Proc Natl Acad Sci U S A 118. 10.1073/pnas.2112267118.

73. Matreyek, K.A., Stephany, J.J., Ahler, E., and Fowler, D.M. (2021). Integrating thousands of PTEN variant activity and abundance measurements reveals variant subgroups and new dominant negatives in cancers. Genome Med 13, 165. 10.1186/s13073-021-00984-x.

74. Papa, A., Wan, L., Bonora, M., Salmena, L., Song, M.S., Hobbs, R.M., Lunardi, A., Webster, K., Ng, C., Newton, R.H., et al. (2014). Cancer-associated PTEN mutants act in a dominant-negative manner to suppress PTEN protein function. Cell 157, 595–610. 10.1016/j.cell.2014.03.027.

75. Ren, P., Xiao, Y., Chang, X., Huang, P.-Y., Li, Z., Gupta, B.B., Chen, X., and Wang, X. (2020). A Survey of Deep Active Learning. arXiv:2009.00236. 10.48550/arXiv.2009.00236.

76. Rao, R., Bhattacharya, N., Thomas, N., Duan, Y., Chen, X., Canny, J., Abbeel, P., and Song, Y.S. (2019). Evaluating Protein Transfer Learning with TAPE. arXiv:1906.08230. 10.48550/arXiv.1906.08230.

77. Shanehsazzadeh, A., Belanger, D., and Dohan, D. (2020). Is Transfer Learning Necessary for Protein Landscape Prediction?, arXiv:2011.03443. 10.48550/arXiv.2011.03443.

78. Xu, M., Zhang, Z., Lu, J., Zhu, Z., Zhang, Y., Ma, C., Liu, R., and Tang, J. (2022). PEER: A Comprehensive and Multi-Task Benchmark for Protein Sequence Understanding. arXiv:2206.02096. 10.48550/arXiv.2206.02096.

79. Ruff, K.M., and Pappu, R.V. (2021). AlphaFold and Implications for Intrinsically Disordered Proteins. J Mol Biol 433, 167208. 10.1016/j.jmb.2021.167208.

80. Muller, H.J. (1932). Further studies on the nature and causes of gene mutations. In Proceedings of the Sixth International Congress on Genetics.

81. Page, D.J., Miossec, M.J., Williams, S.G., Monaghan, R.M., Fotiou, E., Cordell, H.J., Sutcliffe, L., Topf, A., Bourgey, M., Bourque, G., et al. (2019). Whole Exome Sequencing Reveals the Major Genetic Contributors to Nonsyndromic Tetralogy of Fallot. Circ Res 124, 553–563. 10.1161/CIRCRESAHA.118.313250.

82. Thölke, P., and De Fabritiis, G. (2022). TorchMD-NET: Equivariant Transformers for Neural Network based Molecular Potentials. arXiv:2202.02541. 10.48550/arXiv.2202.02541.

83. Kingma, D.P., and Ba, J. (2014). Adam: A Method for Stochastic Optimization. arXiv:1412.6980. 10.48550/arXiv.1412.6980.

84. Liu, X., Li, C., Mou, C., Dong, Y., and Tu, Y. (2020). dbNSFP v4: a comprehensive database of transcript-specific functional predictions and annotations for human nonsynonymous and splice-site SNVs. Genome Med 12, 103. 10.1186/s13073-020-00803-9.

