## Supplementary Figures for "PreMode predicts mode-of-action of missense variants by deep graph representation learning of protein sequence and structural context"

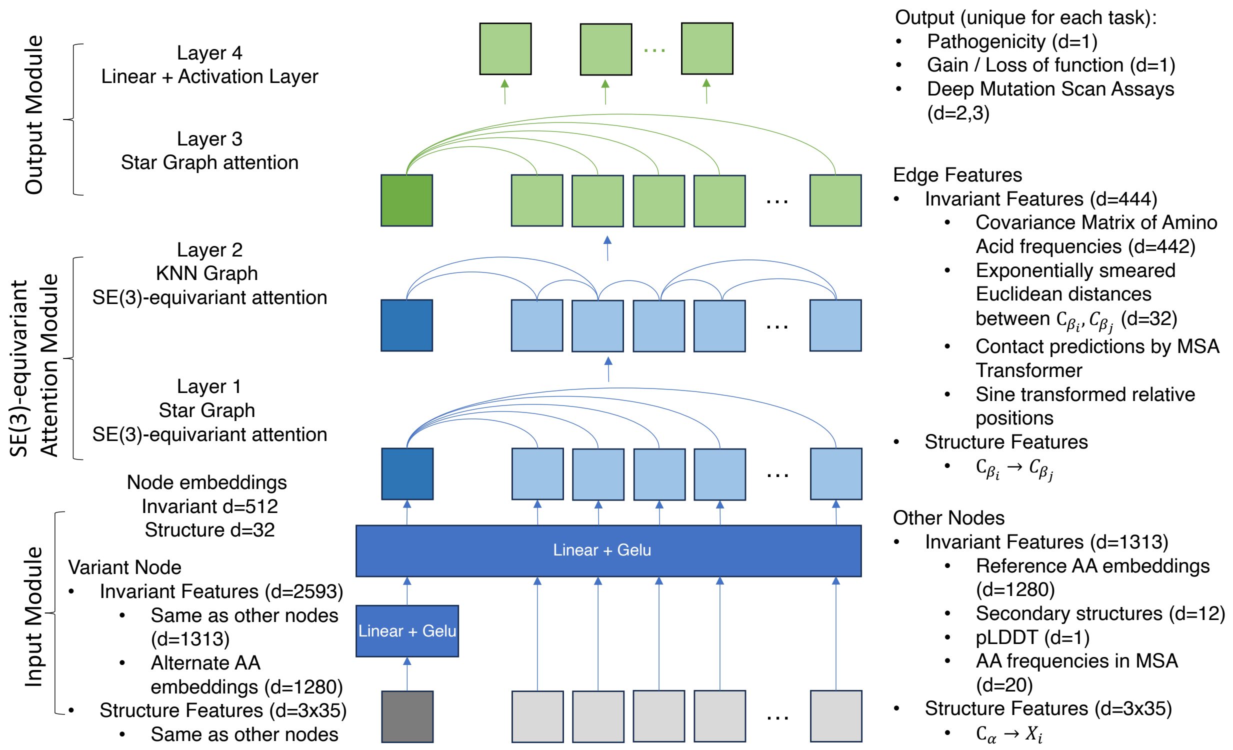


Supplementary Figure 1. Architecture of PreMode. Layers shared during both pretrain and transfer learning were colored blue. Layers only updated during transfer learning were colored green. Arrows indicate data flow.

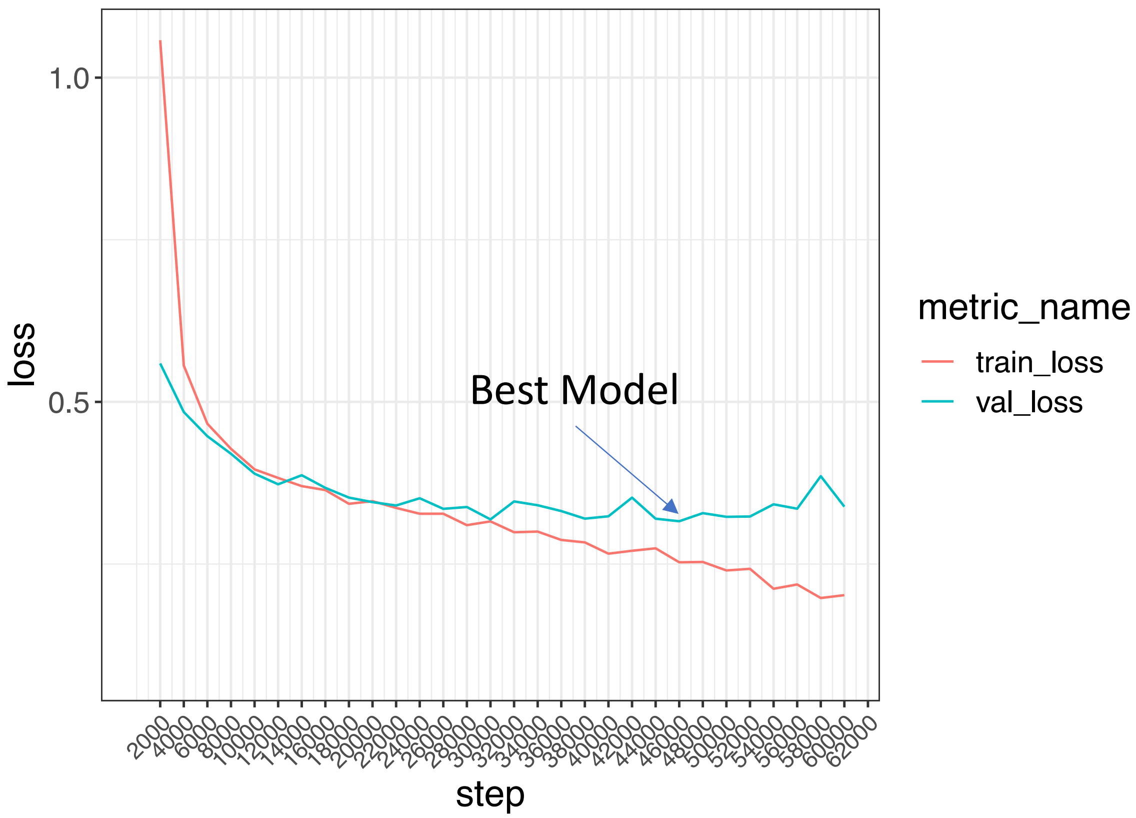


Supplementary Figure 2. Pretrain and model selection of transfer learning starting states. X-axis is training steps, y-axis is loss. Red and blue indicates training and validation loss, respectively.

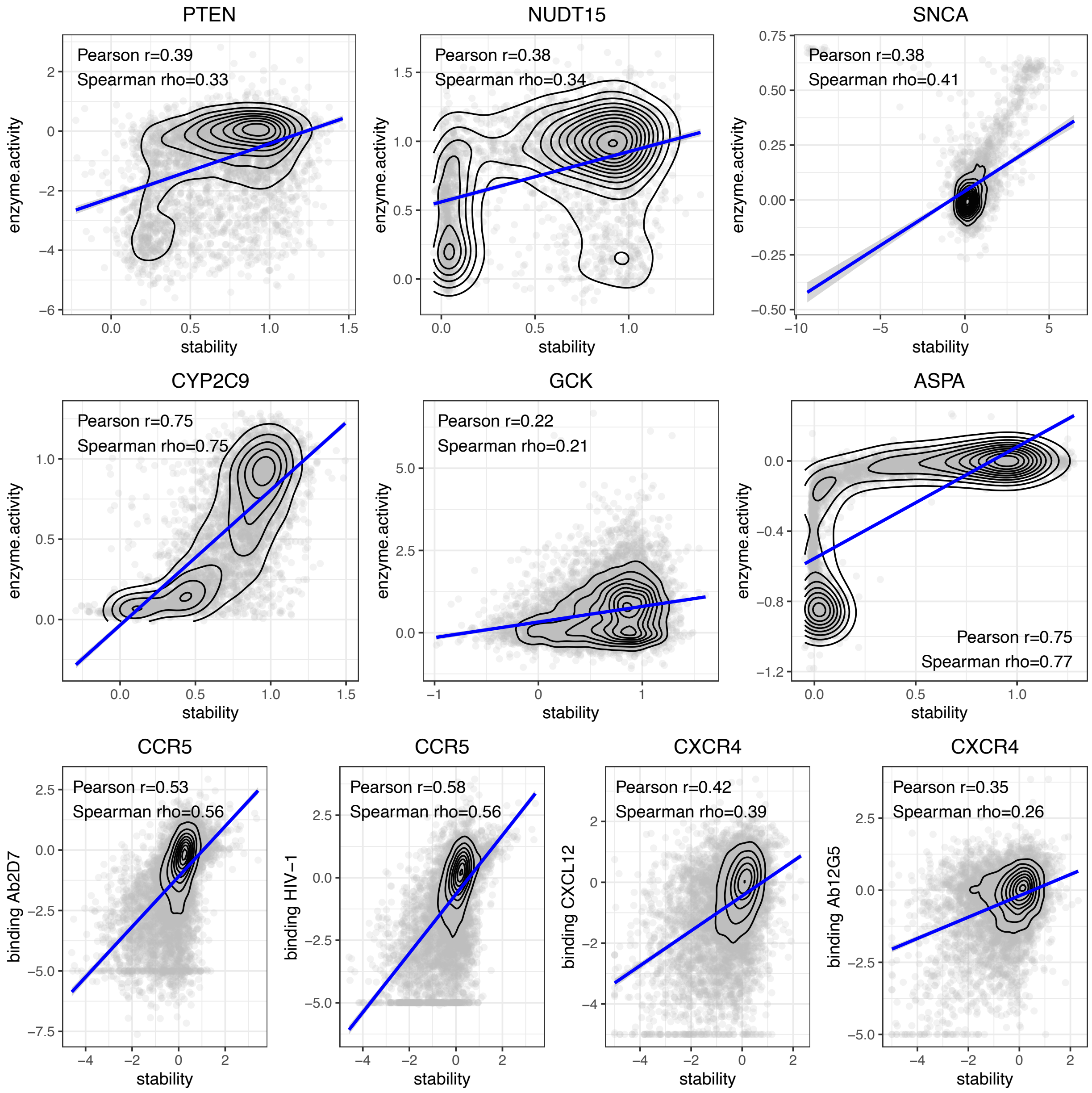


Supplementary Figure 3. Comparison of stability and functional based deep mutational scan experiments. Y-axis, functional-based DMS measurements. X-axis, stability-based measurements.

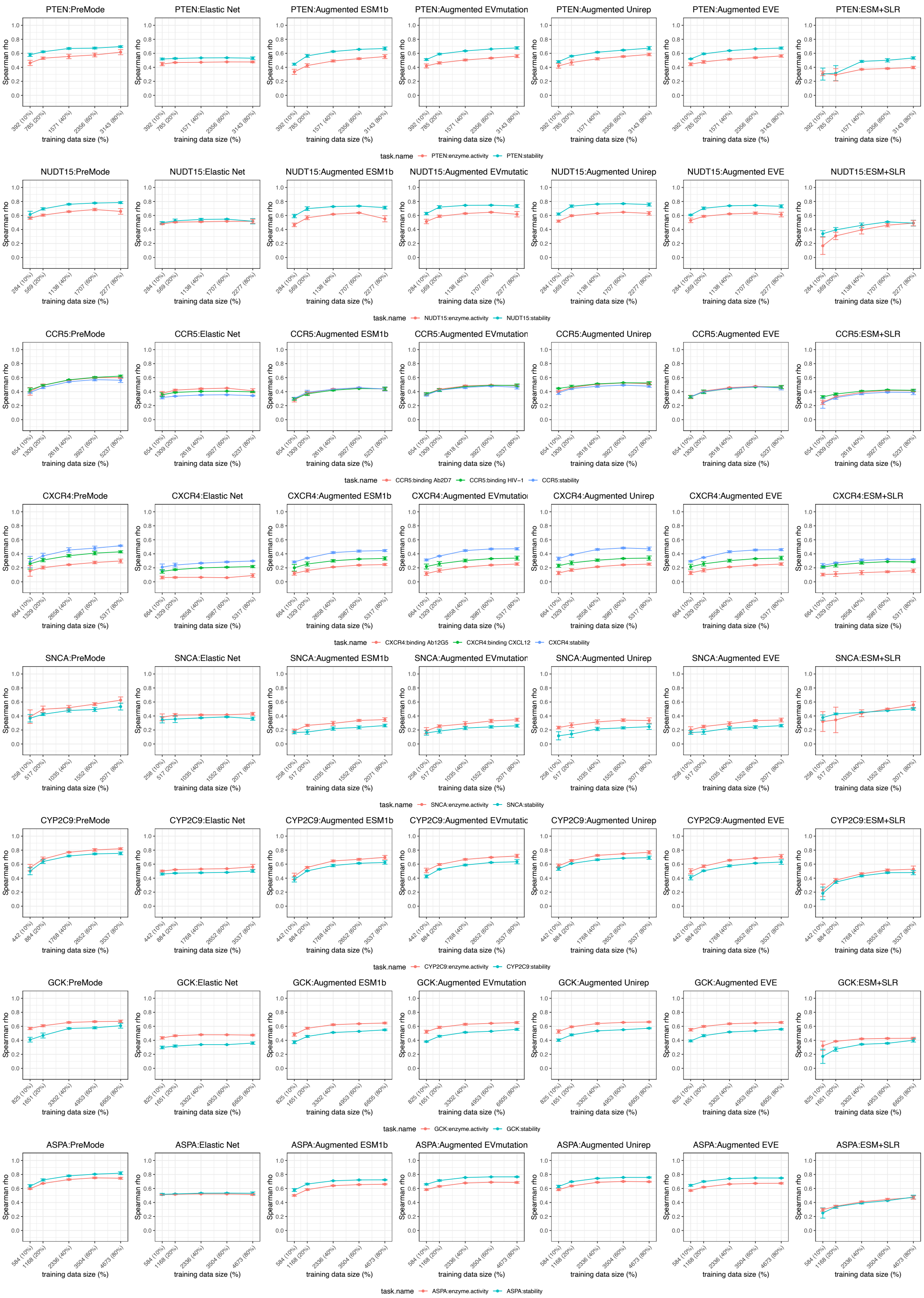
PTEN:PreMode
PTEN:Elastic Net
PTEN:Augmented ESM1b
PTEN:Augmented EVmutation
PTEN:Augmented Unirep
PTEN:Augmented EVE
PTEN:ESM+SLR
Spearman rho
training data size (%)
task.name PTEN:enzyme.activity PTEN:stability
NUDT15:PreMode
NUDT15:Elastic Net
NUDT15:Augmented ESM1b
NUDT15:Augmented EVmutatic
NUDT15:Augmented Unirep
NUDT15:Augmented EVE
NUDT15:ESM+SLR
task.name NUDT15:enzyme.activity NUDT15:stability
CCR5:PreMode
CCR5:Elastic Net
CCR5:Augmented ESM1b
CCR5:Augmented EVmutation
CCR5:Augmented Unirep
CCR5:Augmented EVE
CCR5:ESM+SLR
task.name CCR5:binding Ab2D7 CCR5:binding HIV−1 CCR5:stability
CXCR4:PreMode
CXCR4:Elastic Net
CXCR4:Augmented ESM1b
CXCR4:Augmented EVmutatio
CXCR4:Augmented Unirep
CXCR4:Augmented EVE
CXCR4:ESM+SLR
task.name CXCR4:binding Ab12G5 CXCR4:binding CXCL12 CXCR4:stability
SNCA:PreMode
SNCA:Elastic Net
SNCA:Augmented ESM1b
SNCA:Augmented EVmutation
SNCA:Augmented Unirep
SNCA:Augmented EVE
SNCA:ESM+SLR
task.name SNCA:enzyme.activity SNCA:stability
CYP2C9:PreMode
CYP2C9:Elastic Net
CYP2C9:Augmented ESM1b
CYP2C9:Augmented EVmutation
CYP2C9:Augmented Unirep
CYP2C9:Augmented EVE
CYP2C9:ESM+SLR
task.name CYP2C9:enzyme.activity CYP2C9:stability
GCK:PreMode
GCK:Elastic Net
GCK:Augmented ESM1b
GCK:Augmented EVmutation
GCK:Augmented Unirep
GCK:Augmented EVE
GCK:ESM+SLR
task.name GCK:enzyme.activity GCK:stability
ASPA:PreMode
ASPA:Elastic Net
ASPA:Augmented ESM1b
ASPA:Augmented EVmutation
ASPA:Augmented Unirep
ASPA:Augmented EVE
ASPA:ESM+SLR
task.name ASPA:enzyme.activity ASPA:stability

Supplementary Figure 4. PreMode's transfer learning performances compared to other methods with reduced training data in *PTEN*, *NUDT15*, *CCR5*, *CXCR4*, *SNCA*, *CYP2C9*, *GCK*, *ASPA*. X-axis, amount of data in training. Y-axis, spearman correlation with testing data. Color indicates experimental assays on different protein properties.

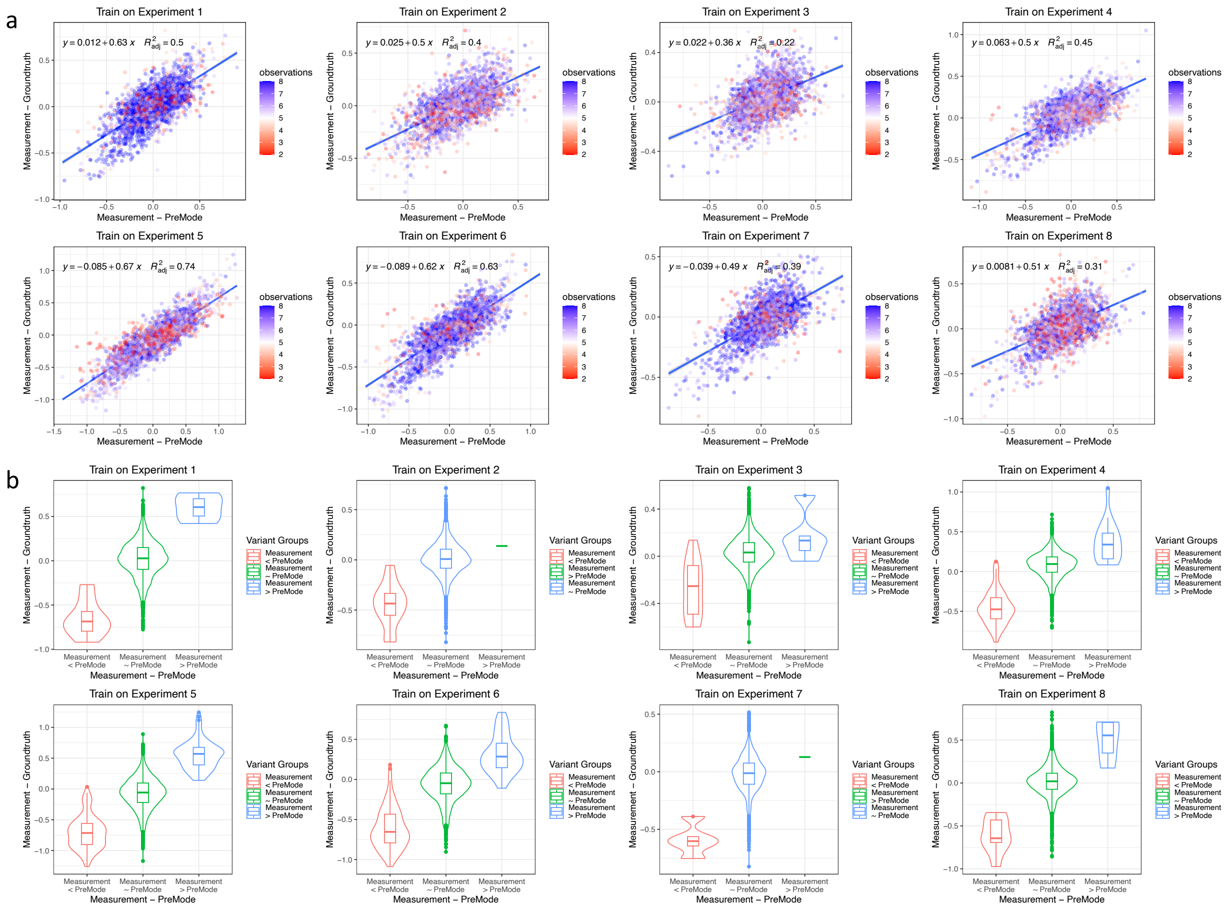


Supplementary Figure 5. PreMode improved PTEN stability deep mutational scan experiments. a) Difference between single experimental readouts and PreMode predictions (X-axis) versus difference between single experimental readouts and average of the rest 7 biological replicates (Y-axis). Points were colored by total number of biological replicates. b) X-axis, difference between single experiment readouts and PreMode predictions, categorized by three groups. “Measurement > PreMode” means measured value larger than PreMode predictions, “Measurement ~ PreMode” means measured value close to PreMode predictions, “Measurement < PreMode” means measured value smaller than PreMode predictions. Y-axis, difference between single experimental readouts and average of the rest 7 biological replicates.

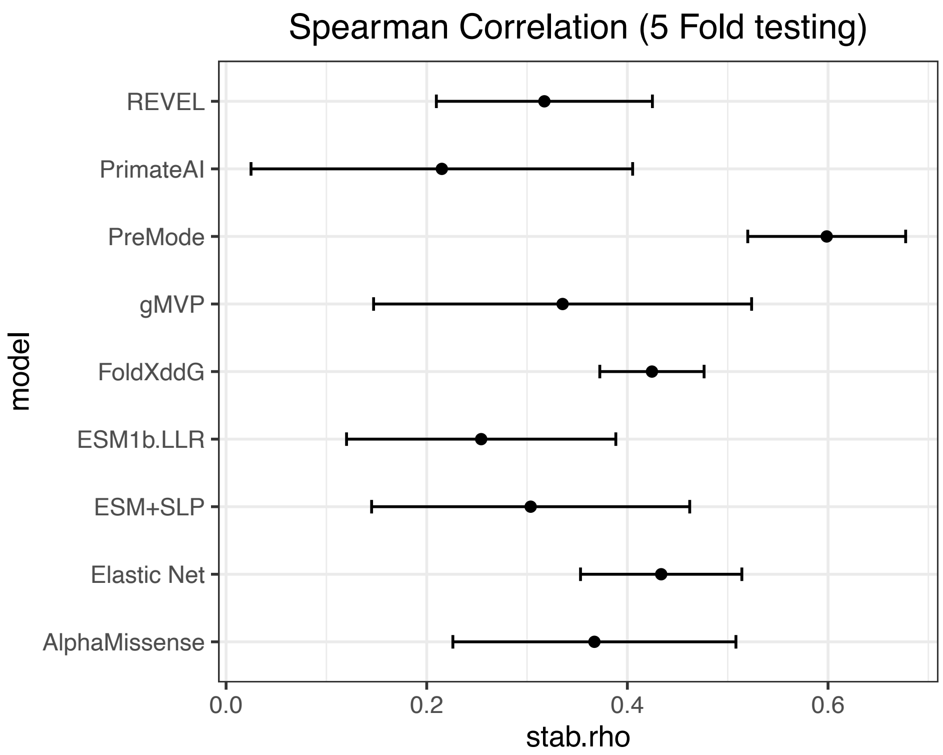


Supplementary Figure 6. Comparison of PreMode and other models on deep mutational scan on stability of untrained genes. Spearman correlations were calculated and averaged by random training/testing splitting 5 times. Error bars show the standard error.

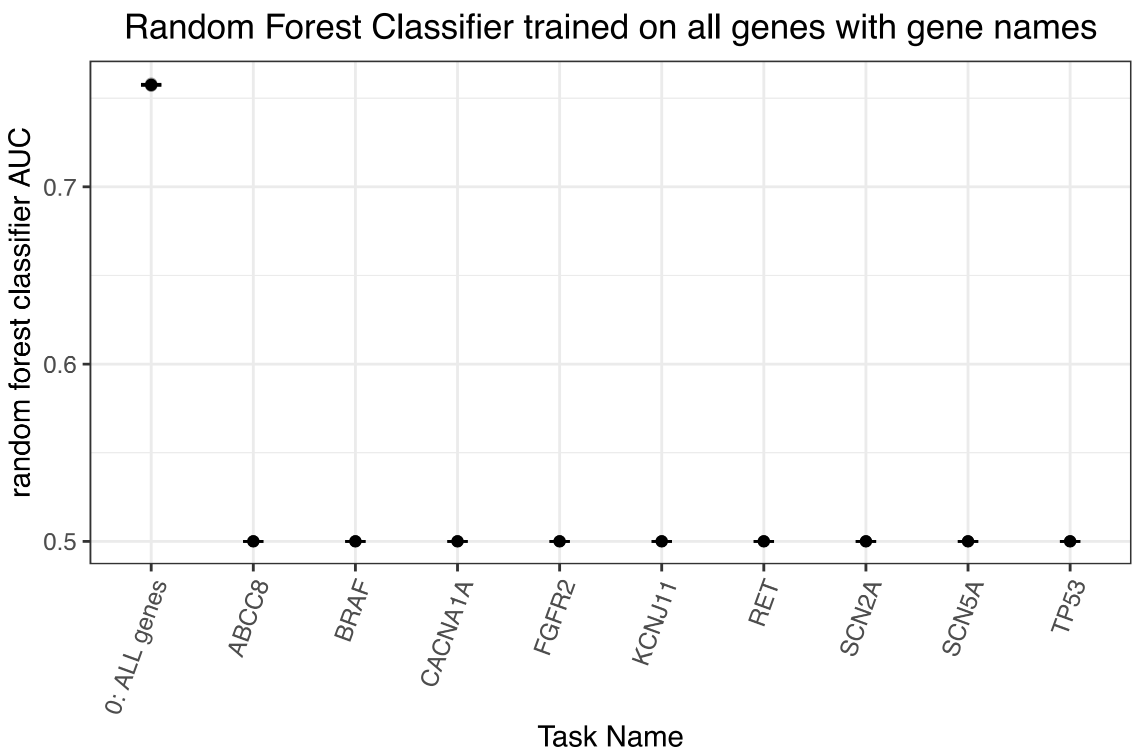


Supplementary Figure 7. Comparison of G/LoF prediction performances using a Random Forest Classifier using gene names as input on whole genome or specific genes. X-axis, gene names, y-axis, AUROC calculated and averaged by random training/testing splitting 5 times, error bars show the standard error.

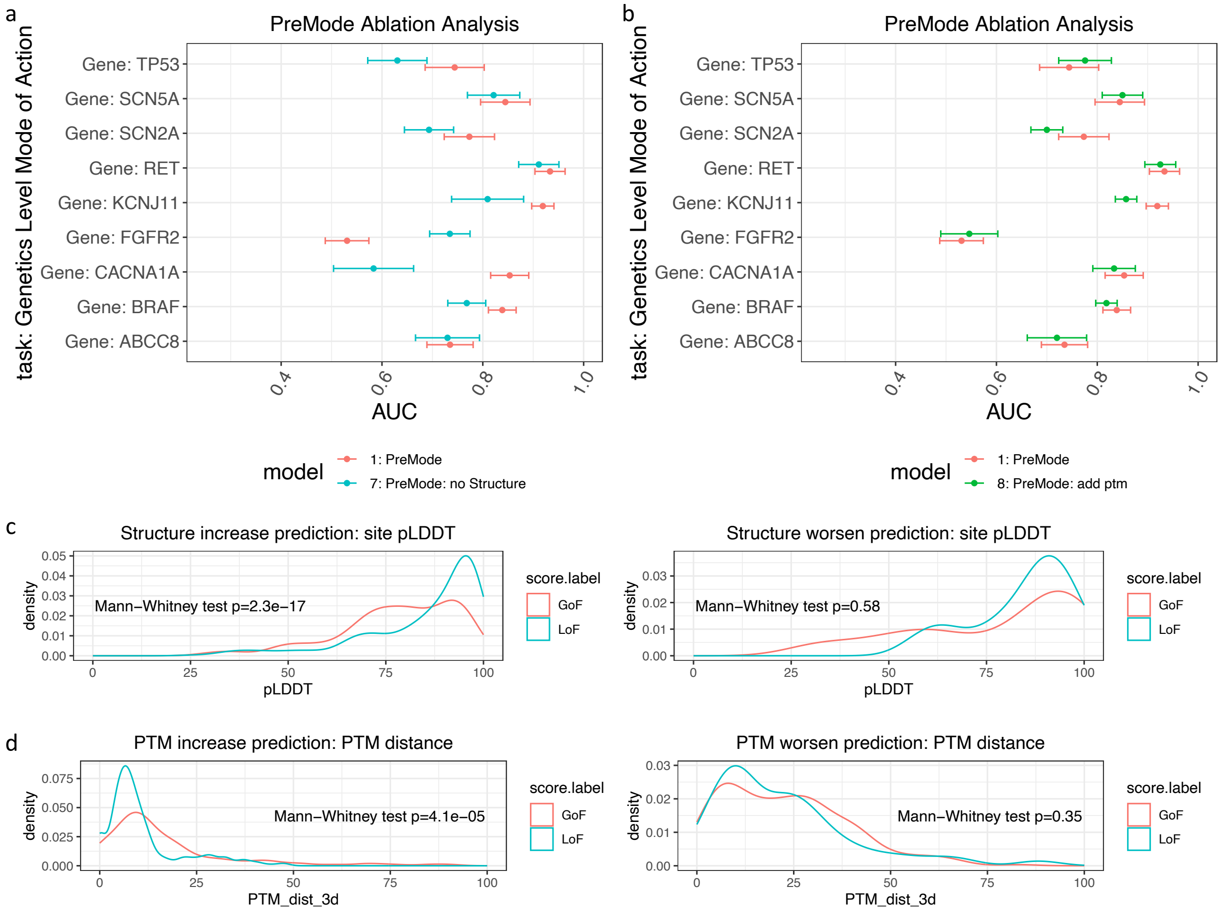


Supplementary Figure 8. A) Comparison of PreMode and PreMode trained without structure information on nine genes, error bars show the standard error. B) Comparison of PreMode and PreMode trained using one hot-encoded PTM as additional input on nine genes, error bars show the standard error. C) Comparison of variant site pLDDT on *FGFR2* (right) versus other eight genes (left), colored by gain or loss-of-function variants. D) Comparison of distances between variant site to the closest PTM site on *TP53, SCN5A* and *FGFR2* (left) versus other five genes (right), colored by gain or loss-of-function variants.

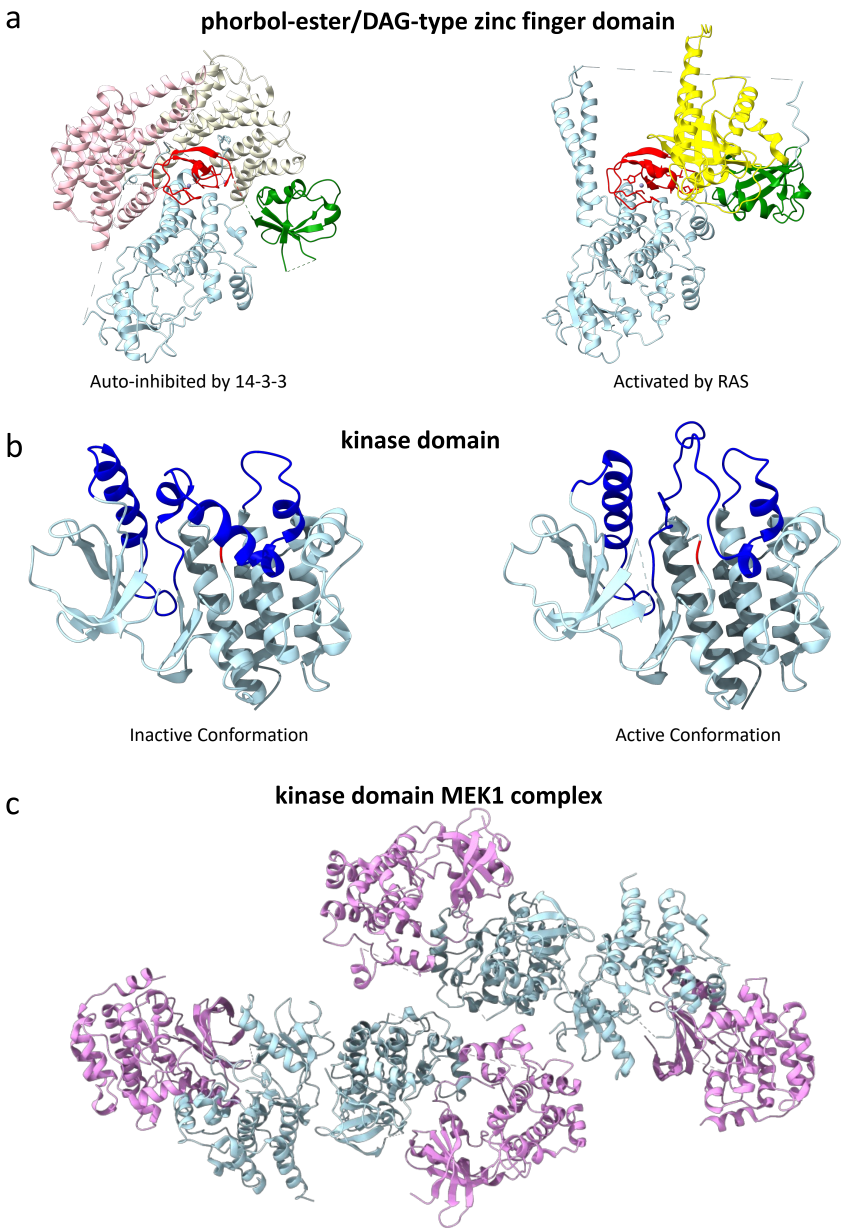


Supplementary Figure 9. Braf functional structures. A) Braf binding to 14-3-3 (left, PDB: 7MFD) and Kras (right, ColabFold predicted) structures. Red is Braf phorbol-ester/DAG-type zinc finger domain, green is Braf Ras binding domain (RBD), light blue is the rest of Braf, pink and white are two 14-3-3 proteins, yellow is Kras. B) Braf kinase domain inactive state conformation (left panel, PDB: 4EHE) and active state conformation (right panel, PDB: 4MNE), red is the active site, blue shows the flexible region, light blue shows the rest of kinase domain. C) Structure of BRAF-MEK1 complex (PDB: 4MNE). BRAF chains were colored light blue, MEK1 chains were colored pink.

a

b

c

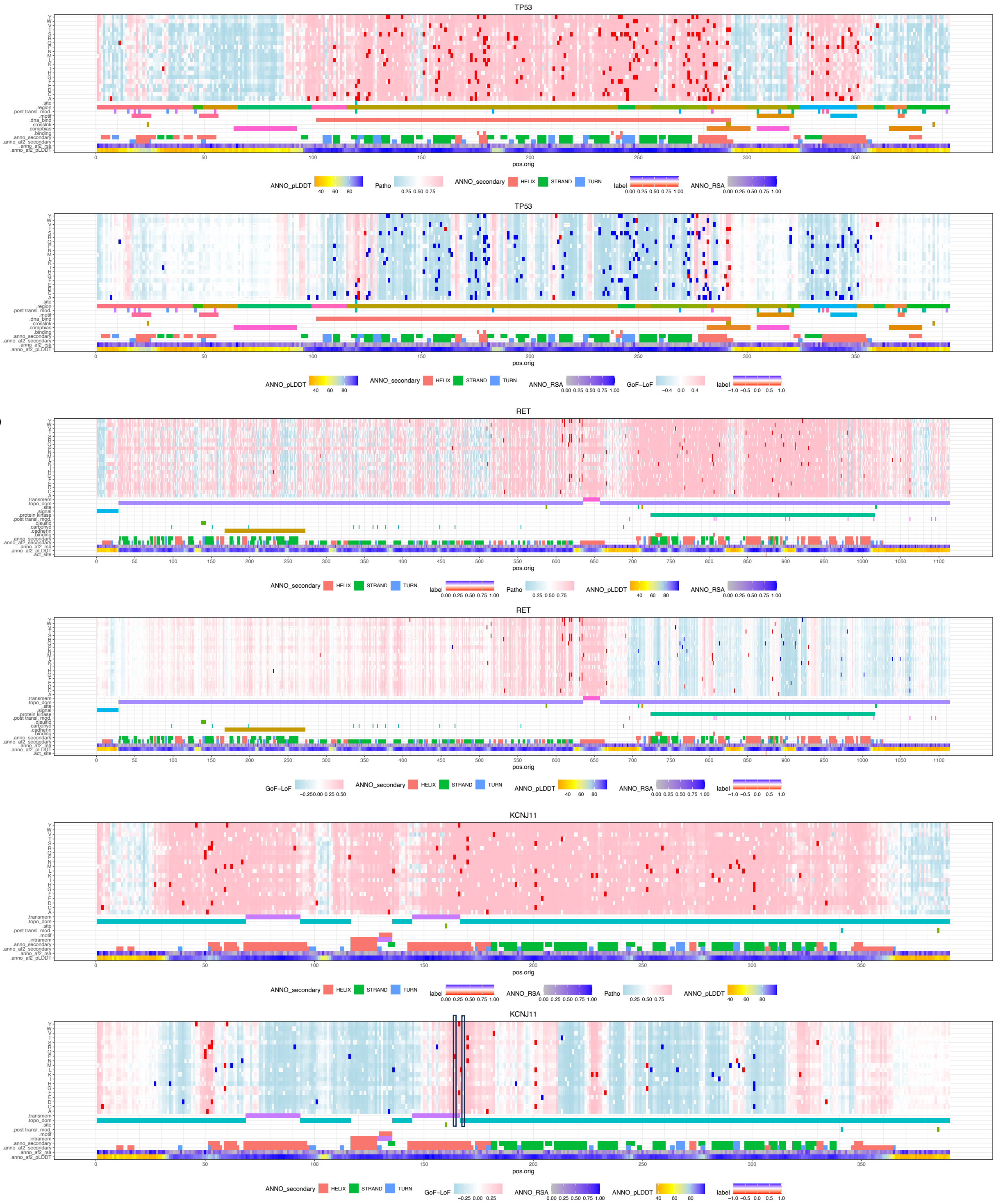


Supplementary Figure 10. In silico mutagenesis results of TP53, RET and KCNJ11. X-axis is amino acid positions from N-terminal to C-terminal. Y-axis shows the amino acid changes, protein domain annotations, Alphafold2 pLDDT, secondary structures and relevant solvent accessibility. The upper panel shows the PreMode pathogenicity prediction, where pink, light blue, red and blue indicates predicted pathogenic, predicted benign, labeled pathogenic, labeled blue, respectively. The lower panel shows the PreMode G/LoF prediction, where pink, light blue, white, red, and blue indicates predicted GoF, predicted LoF, predicted benign, labeled GoF, labeled LoF, respectively.

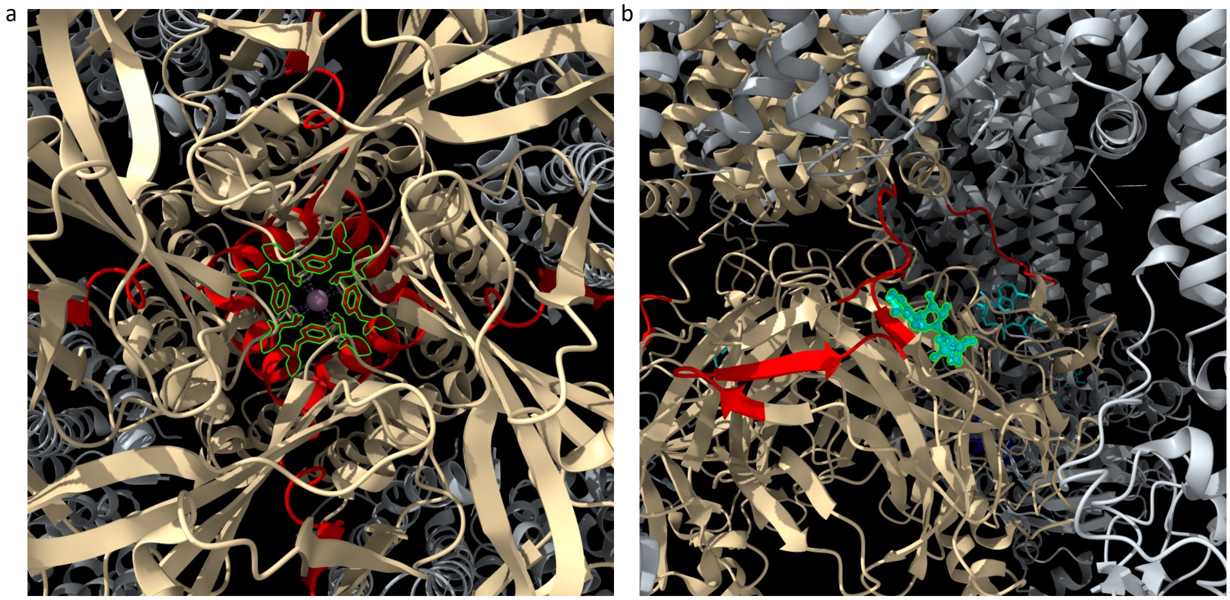

Supplementary Figure 11. Structural view of the KCNJ11 GoF variants enriched domains. A) Cytoplasmic side view of the GoF variants enriched domain (160-190) in KCNJ11 (PDB: 6C3O). Four identical chains of KCNJ11 and ABCC8 were colored as light yellow, grey respectively. The potassium ion was colored as purple. The GoF enriched domains (160-190) were colored as red. L164 and F168 were highlighted as green. B) Side view of the GoF variants enriched domains (red, 44-54, 179-185, 328-340) in KCNJ11 (PDB: 6C3O). ATP was colored as cyan.

a

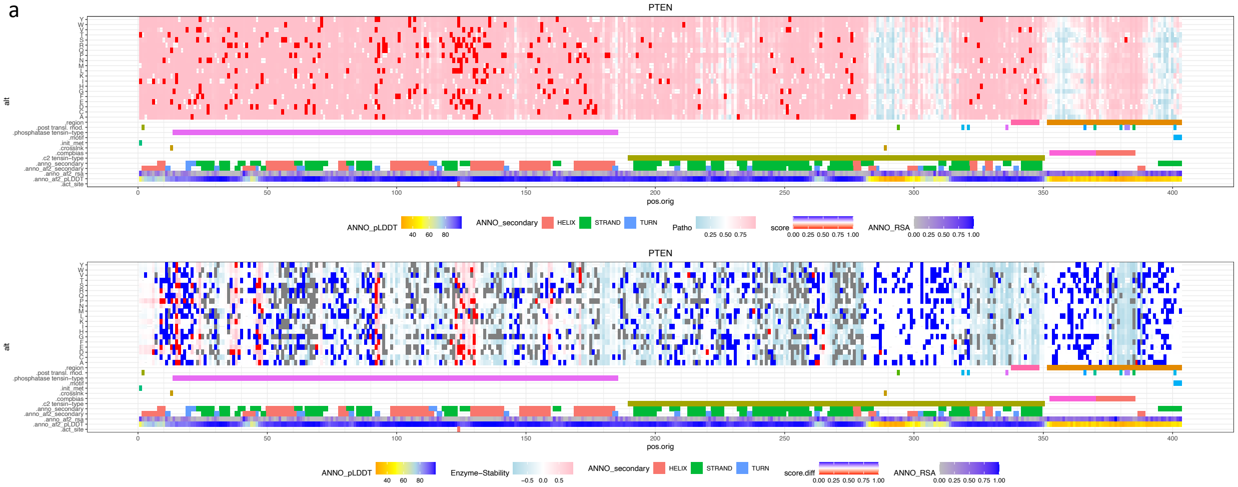


b

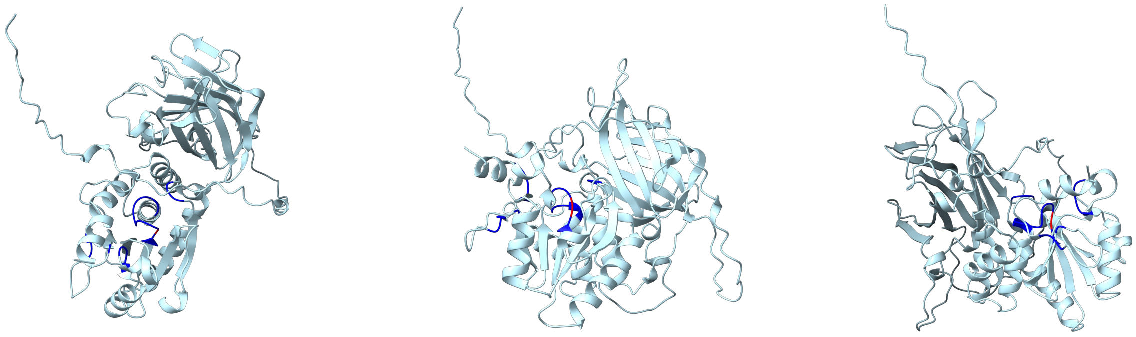

Top View Right View Left View

Supplementary Figure 12. Different loss of function variants in PTEN identified by experiments and predicted by PreMode. a) X-axis is amino acid positions from N-terminal to C-terminal. Y-axis shows the amino acid changes, protein domain annotations, Alphafold2 pLDDT, secondary structures and relevant solvent accessibility. The upper panel shows the PreMode pathogenicity prediction, where pink, light blue, red and blue indicates predicted pathogenic, predicted benign, labeled pathogenic, labeled blue, respectively. The lower panel shows the PreMode different LoF prediction, where pink, light blue, red, blue, and grey indicates predicted function loss only, predicted stability loss only, labeled function loss only, labeled stability loss only, labeled both stability and function loss, respectively. b) Protein structure of PTEN. Blue indicates regions enriched of function loss only variants, red is the PTEN active site.

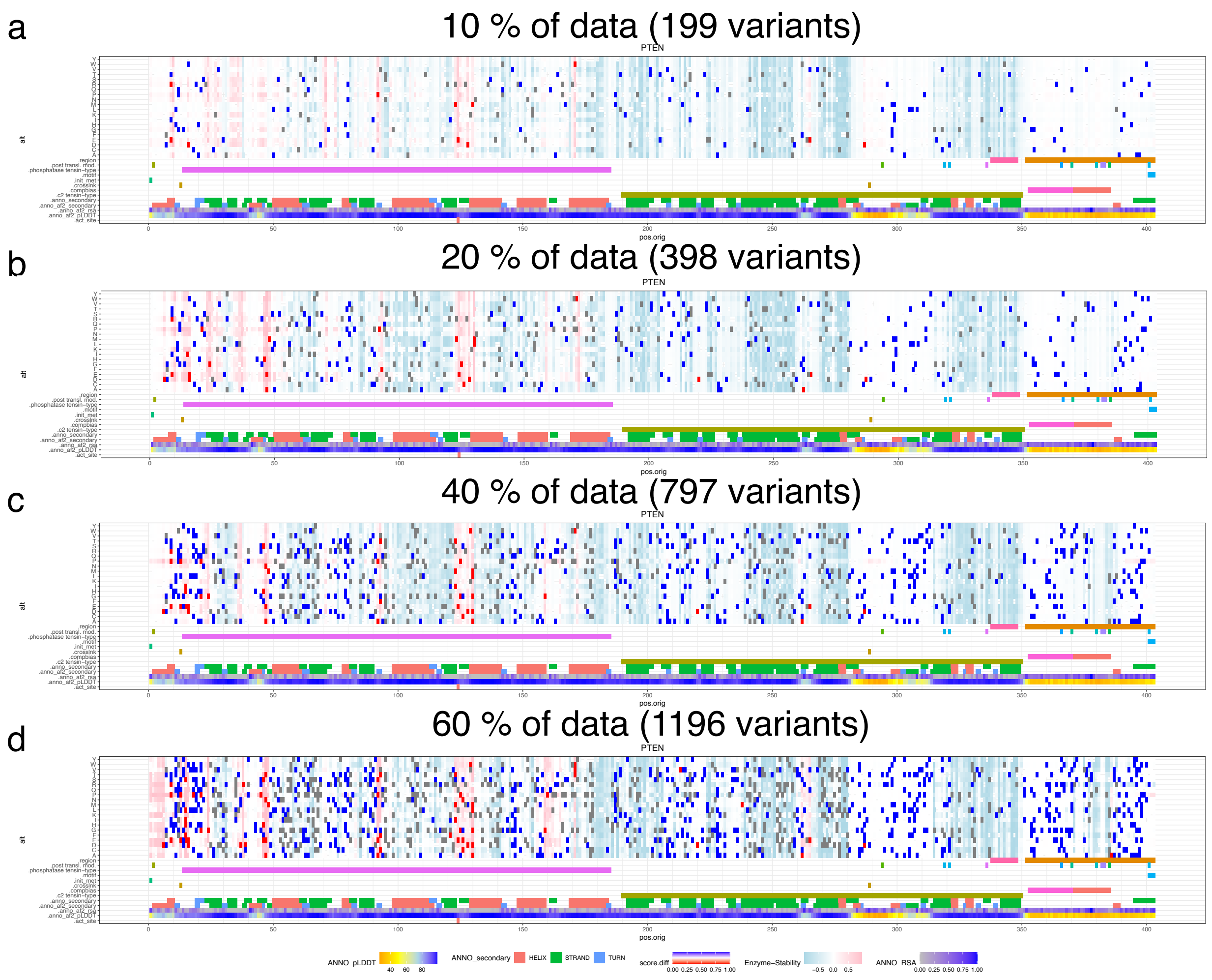


Supplementary Figure 13. PreMode's performance in identifying the enzyme pocket enriched of variants that disrupt enzyme function but maintain protein stability with less training data, colored in same scheme as Supplementary Figure 12. a) 10% of training data. b) 20% of data. c) 40% of data. d) 60% of data.

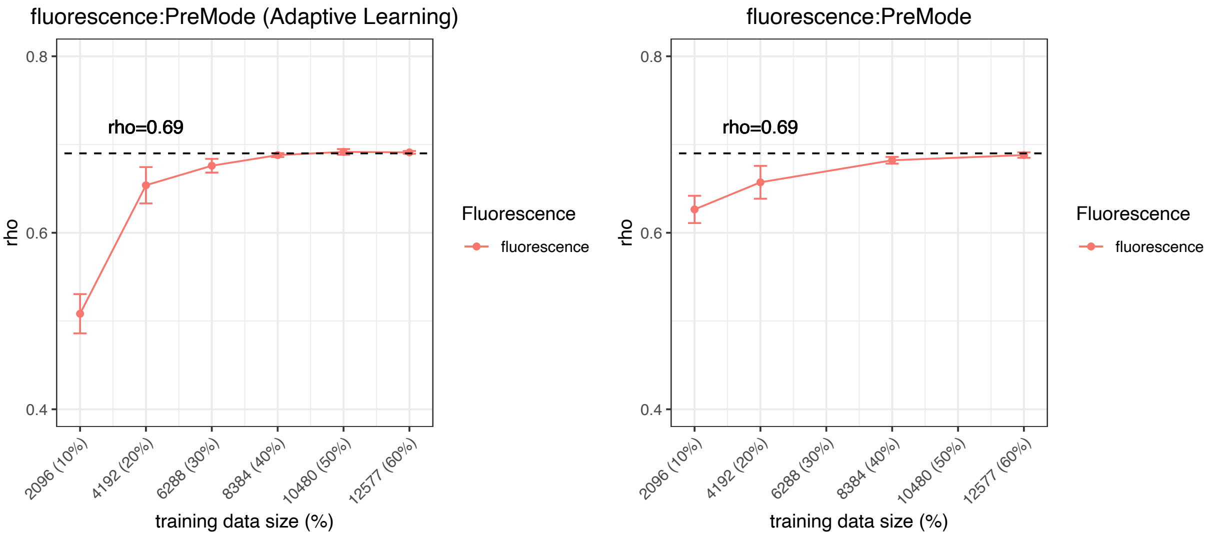


Supplementary Figure 14. PreMode's performance in active learning application to green fluorescence protein (GFP) stability prediction. Left panel shows the performance of active learning with regards to training sample sizes. Right panel shows performances without active learning. Dash line shows the state-of-the-art model currently available in literature.
